# Accurate and interpretable protein complex stoichiometry prediction through agentic AI with StoiAgent

**DOI:** 10.64898/2026.11.17.767325

**Authors:** Jian Liu, Pawan Neupane, Jianlin Cheng

## Abstract

Accurate protein complex structure modeling requires knowledge of complex stoichiometry, yet the copy numbers of individual subunits are often unknown. Consequently, predicting protein complex stoichiometry has emerged as a major challenge in recent years. Here, we present StoiAgent, a large language model (LLM)-powered AI agent that automates the retrieval and integration of sequence, structural, and literature evidence to predict stoichiometry while preserving interpretable reasoning traces (chain of thoughts) of its evidence assessment and ranking decision. An open-weight LLM model, Qwen3.6-27B, is used by StoiAgent as the LLM backbone to combine sequences, structural templates, AlphaFold3 structure predictions, and literature evidence, and perform stoichiometry candidate generation and ranking. On 56 CASP17 protein complex targets, it achieves top-1 accuracy of 75.0%, which represents a 23.2 percentage-point improvement over PreStoi with AlphaFold3 ranking (51.8%) and matches the top-1 accuracy of StoPred. Moreover, StoiAgent has a higher top-3 and top-5 accuracy than StoPred (85.7% versus 80.4% and 87.5% versus 85.7%, respectively). Furthermore, it outperforms general-purpose frontier LLMs, including ChatGPT and Gemini, in stoichiometry prediction. Case studies further illustrate how StoiAgent evaluates competing assembly hypotheses by integrating complementary evidence, with recorded reasoning traces documenting its decision-making process. Overall, StoiAgent demonstrates the potential of LLM-powered AI agents to improve protein complex stoichiometry prediction through the automated integration of diverse sources of evidence while providing interpretable explanations to guide downstream structure modeling.

## Introduction

Proteins carry out many essential cellular functions through interactions with other proteins, forming complexes that participate in processes such as signal transduction, enzymatic catalysis, molecular transport, and gene regulation. These complexes range from simple dimers to large molecular assemblies composed of many subunits (chains). Their biological functions depend not only on the identities and three-dimensional structures of their constituent proteins but also on how these proteins assemble and interact [1].

Recent advances in deep learning have substantially improved the accuracy of protein complex structure prediction. However, widely used methods, including AlphaFold3 [2], Chai-1 [3], and Boltz-2 [4], require the composition information of a target complex (stoichiometry), i.e., the number of copies of each unique protein subunit, as input to predict its structure. This requirement presents a challenge because the stoichiometry of most protein complexes is unknown. Moreover, different stoichiometries involving the same proteins can result in different assembly sizes, symmetries, interaction interfaces, and biological functions [5]. Therefore, it is important to predict the stoichiometry of protein complexes, providing essential information for accurate complex structure prediction.

Predicting protein complex stoichiometry remains challenging because the amino acid sequences of individual proteins do not directly specify how many copies of each subunit assemble into a functional complex. Homology-based approaches can provide valuable clues by identifying related, experimentally characterized complexes with known stoichiometries [6] as templates. Highly significant sequence matches to structural templates can provide strong evidence for a candidate assembly when the relevant oligomeric organization is conserved. However, many protein complexes do not have similar templates. Even when some templates are available, the reliability of the evidence still depends on the relationship between the target and template complexes, the completeness of the recorded assembly, and the conservation of oligomeric state [7]. Moreover, different templates may support different candidate stoichiometries, making it difficult to identify the correct one. Structure prediction methods can help evaluate these competing candidates by assessing their structural plausibility. However, high prediction confidence does not necessarily establish the biological stoichiometry [8]. These limitations motivate the development of methods that integrate sequence, template, and structural evidence to systematically evaluate and rank candidate stoichiometries, thereby improving the identification and modeling of biologically relevant protein assemblies.

Two recent sequence-based deep learning approaches have demonstrated promising performance in stoichiometry prediction. StoPred [9] uses protein language model (PLM) representations and graph attention networks to infer copy numbers for each subunit [9], while Stoic incorporates interface-aware representations and offers residue-level interpretability [10]. Although these methods have advanced predictive performance, understanding the evidence supporting a particular stoichiometry and distinguishing among competing assembly hypotheses remain important challenges.

Our previous method, PreStoi, infers and ranks candidate stoichiometries using homologous structural template and literature evidence first and subsequently evaluates candidates using AlphaFold3 ranking scores [11]. However, integrating evidence from different sources and resolving conflicting predictions still requires human judgment, limiting the automation and scalability of the approach. Recent advances in large language models (LLMs) and agentic AI provide an opportunity to automate this process while expanding the evidence available for stoichiometry prediction. Building on PreStoi, we develop StoiAgent, an LLM-powered AI agent that autonomously retrieves external evidence such as literature information and integrates it with sequence, template, and structural information to evaluate and prioritize candidate stoichiometries. Specifically, StoiAgent operates in two stages: candidate proposal and prioritization in Stage 1, followed by structural-score-informed reranking of a fixed shortlist of candidates in Stage 2. By combining automated evidence retrieval and integration, StoiAgent seeks to improve prediction accuracy while preserving reasoning traces (chain of thoughts) that allow researchers to examine the evidence considered in its predictions.

We develop and test StoiAgent for protein complex stoichiometry prediction using the protein complex targets in the 16th and 17th Critical Assessment of Protein Structure Prediction (CASP16 [12] and CASP17). CASP16 serves as the development benchmark for optimizing evidence configurations and selecting the LLM backbone, while CASP17 provides a separate blind benchmark for evaluating StoiAgent against existing stoichiometry prediction methods, including PreStoi, StoPred, and Stoic, as well as general-purpose LLMs such as ChatGPT [13] and Gemini [14]. We further investigate the contributions of LLM-based candidate prioritization, structural-score-informed reranking, and different sources of evidence to prediction performance. Through these evaluations, we demonstrate the potential of LLM-powered agents to improve stoichiometry prediction by automating evidence retrieval and integration while providing interpretable support for predicted assemblies. The overall StoiAgent workflow is illustrated in Fig. 1.

**Figure 1:**
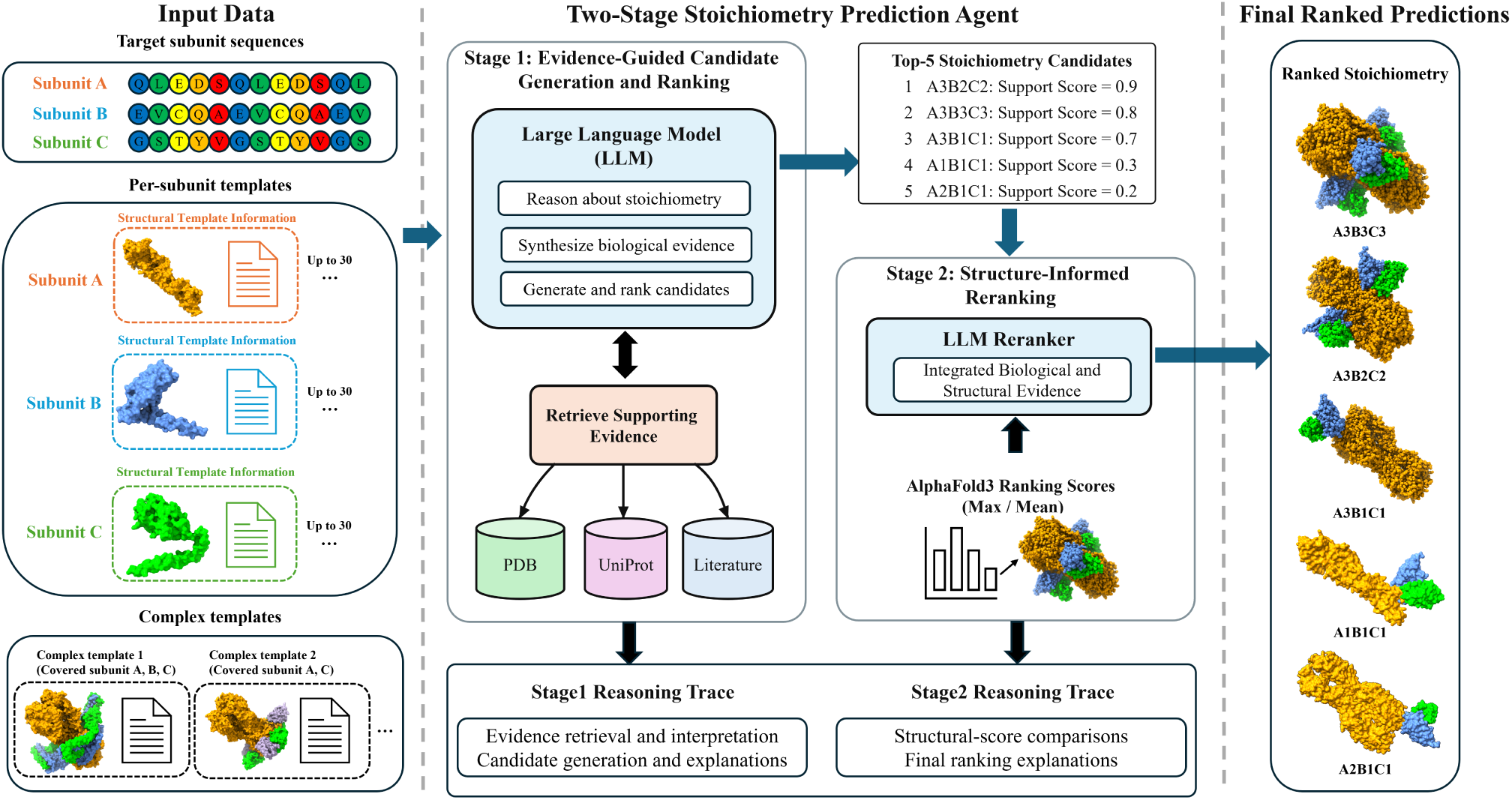
Overview of the StoiAgent workflow. Given protein sequences, target descriptions, and structural-template information, In Stage 1, StoiAgent uses an LLM to retrieve supporting biological evidence from PDB, UniProt, and the literature, integrate the available evidence, and generate and rank candidate stoichiometries with support scores. The top-5 candidates are passed to Stage 2, where the LLM integrates AlphaFold3 ranking scores with Stage-1 evidence to rerank the candidates and produce the final ranked predictions. Both stages preserve reasoning traces documenting evidence assessments and candidate-ranking decisions.

## Results

### Development and evaluation datasets

A total of 28 CASP16 Phase-0 complex targets were used for method development, including 19 heteromeric and 9 homomeric complexes. These targets were used to compare evidence configurations and select the primary LLM backbone. An independent set of 56 CASP17 targets was used to evaluate the selected StoiAgent configuration against existing methods and assess its sensitivity to different LLM backbones.

The CASP17 evaluation dataset comprises 56 targets whose stoichiometries were initially unknown and later revealed for performance evaluation (Fig. 2). Among these targets, 43 are heteromeric and 13 are homomeric, representing 18 distinct native stoichiometries (Fig. 2a). The five most frequent compositions are A1B1C1 (17 targets), A3B3C3 (7), A3 (6), A2 (5), and A2B2C2 (4), collectively accounting for 39 of the 56 targets (69.6%). Less frequent compositions include A8B4, A9, A17, and a 36-chain assembly A3B3C6D18E3F3. To assess how frequently simple stoichiometric assumptions could recover the reference compositions, we considered a minimal-copy baseline that predicts A2 for homomers and one copy of each entity for heteromers. This baseline correctly predicts 30 of the 56 targets (53.6%). These results highlight the importance of evaluating alternative assemblies beyond simple minimal-copy compositions.

**Figure 2:**
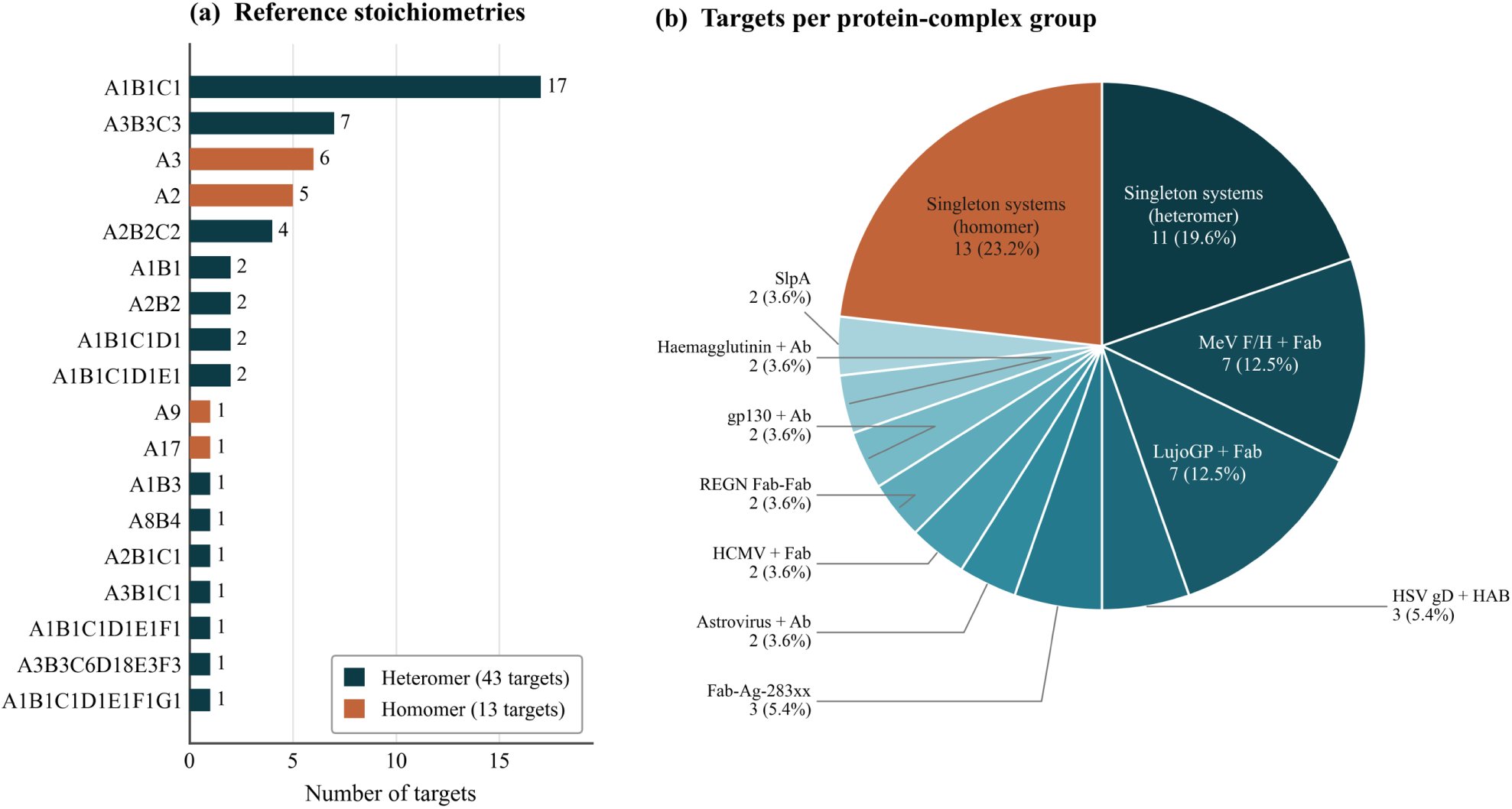
Composition of the CASP17 evaluation dataset. (a) Distribution of 18 native stoichiometries across 43 heteromeric and 13 homomeric targets. (b) Distribution of 56 targets across biological systems. Singleton heteromeric and homomeric systems contribute 11 and 13 targets, respectively, while 10 recurring systems contribute 32 targets. Percentages are calculated relative to all 56 targets. The dataset represents 34 distinct biological systems.

In addition to the variation in stoichiometric composition, the CASP17 targets are unevenly distributed across biological systems (Fig. 2b). The 56 targets represent 34 annotated biological systems, of which 24 contribute one target each, while the remaining 10 contribute 32 targets. In particular, MeV F/H + Fab and LujoGP + Fab complexes each contribute 7 targets. All 13 homomeric targets represent distinct biological systems, whereas the 43 heteromeric targets belong to 21 systems. Consequently, conventional target-level accuracy gives greater weight to biological systems represented by multiple targets, potentially influencing comparisons among prediction methods. To account for this imbalance, we additionally report group-balanced performance, which assigns equal weight to each biological system.

To approximate the blind prediction setting of CASP17, we restricted StoiAgent to external evidence available before each target’s initial release date, when its native stoichiometry had not yet been disclosed. Database and literature retrieval were subject to target-specific temporal cutoffs, while structural-template evidence was prepared using a template database curated from PDB [15] with a fixed cutoff of April 1, 2026 (prior to the start date of CASP17). These restrictions were applied across the evaluated LLM backbones and evidence configurations. Further implementation details are provided in the Methods section.

### Selection of LLM backbone and input configuration using CASP16 dataset

We evaluate four open-weight LLM backbones, including Qwen3.6-27B [16], Gemma-3-27B-it [17], OLMo-2-32B-Instruct [18] and Mistral-Small-3.2-24B [19], on 28 CASP16 targets using the same candidate-generation procedure and inference settings (Fig. 3) in terms of top-1 accuracy, top-3 accuracy, and top-5 accuracy. Top-1 accuracy is the percentage of correct stoichiometry predictions that are correct when only one stoichiometry is predicted for each target. Top-3 (top-5) accuracy considers top three (or five) predicted stoichiometries for each target, and the prediction is correct if any of the top three (or five) stoichiometries is correct.

**Figure 3:**
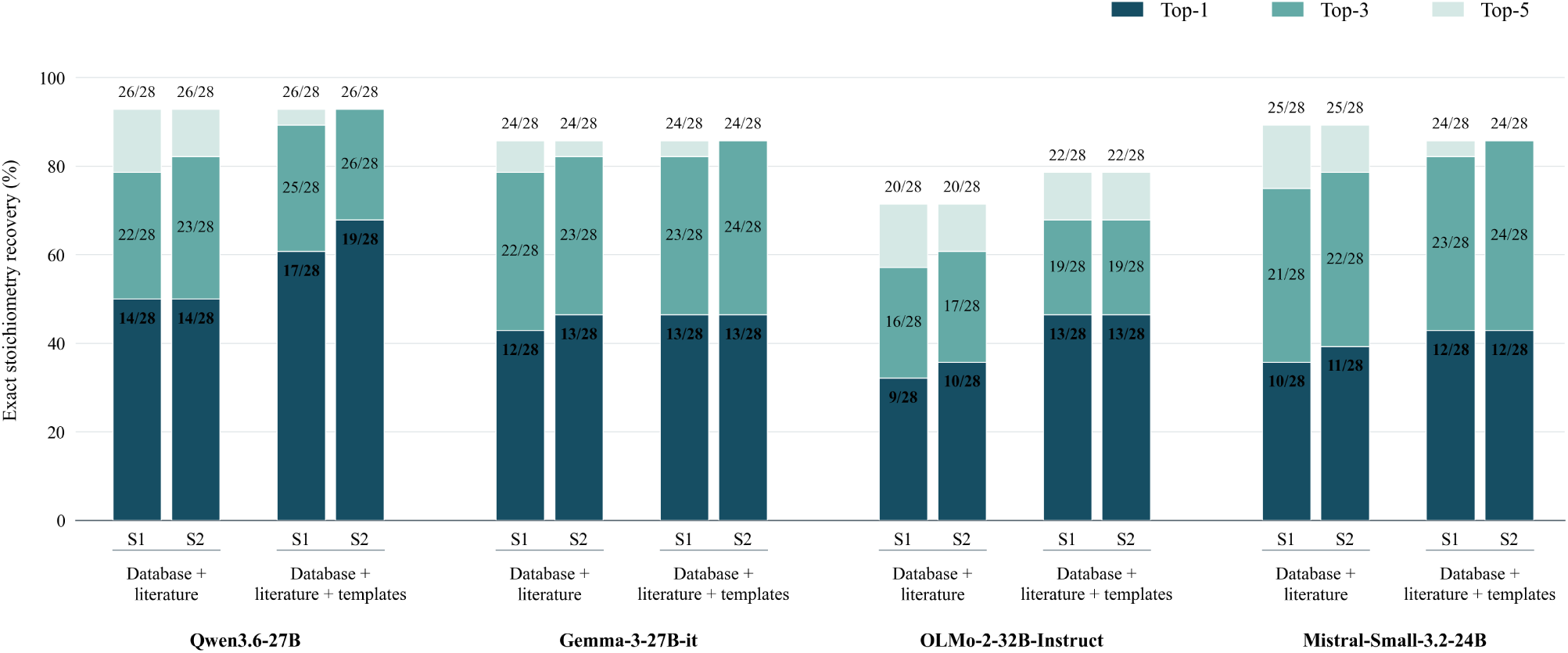
Comparison of four LLM backbones and two evidence configurations on 28 CASP16 targets (19 heteromers and 9 homomers). Each LLM is evaluated using two input configurations: database and literature retrieval, with or without additional structural template evidence. Stage 1 (S1) generates and ranks candidate stoichiometries, while Stage 2 (S2) reranks the fixed top-5 candidates considering AlphaFold3 ranking scores. Stacked bars indicate cumulative top-1 accuracy (dark blue), top-3 accuracy (blue), and top-5 accuracy (light blue). Numeric labels indicate the corresponding numbers of correctly predicted targets. Missing or invalid predictions are counted as incorrect.

Two input evidence configurations are compared: Database + literature and Database + literature + templates. Both configurations provide the amino acid sequences of individual protein subunits and target descriptions, while allowing the LLMs to retrieve additional biological information from PDB, UniProt [20], and PubMed [21]. The second configuration additionally supplied structural template evidence obtained from sequence-based template searches against known structures in the PDB. For heteromeric complexes, templates matching at least two distinct target subunits are prioritized to provide evidence of their potential assembly organization. When no eligible shared template is available, individual-subunit templates are used instead. For homomeric complexes, individual-subunit templates with relevant information such as PDB code, e-value and copy number are provided. Each LLM and evidence configuration is evaluated in both Stage-1 candidate generation and Stage-2 AlphaFold3-score-informed reranking.

As shown in Fig. 3, Qwen3.6-27B achieves the better prediction accuracy than the other three LLMs in almost all situations, except in two cases it has the same top-3 accuracy as Gemma-3-27B-it. Moreover, using Database + Literature + Template generally yields better accuracy than using Database + Literature or at least the same accuracy for all four LLMs backbones. Furthermore, considering AlphaFold3 ranking scores in stoichiometry reranking in Stage 2 generally improves the performance over Stage 1 without the reranking.

Particularly, Qwen3.6-27B with Database + Literature + Template input achieves the highest top-1 accuracy in Stage 2: 19/28 (67.9%), higher than its accuracy of 17/28 (60.7%) using the same input configuration in Stage 1 and 14/28 (50.0%) using Database + Literature in Stages 1 and 2. The top-3 and top-5 accuracy of Qwen3.6-27B with Database + Literature + Template input reaches 26/28 (92.9%). Based on the results, we select Qwen3.6-27B with the Database + literature + templates input configuration and the Stage-2 reranking procedure for the subsequent blind evaluation on CASP17 targets.

### StoiAgent outperforms existing stoichiometry prediction methods, ChatGPT, and Gemini on CASP17 dataset

We evaluate StoiAgent on 56 CASP17 targets in comparison with three existing methods, including PreStoi, StoPred, Stoic, and a minimal-copy baseline (Fig. 4A). StoiAgent (Stage 2) achieves a top-1 accuracy of 75.0% (42/56), substantially exceeding that of our previous method, PreStoi (51.8%, 29/56), by 23.2 percentage points. Its top-1 accuracy matches StoPred (75.0%) and is much higher than Stoic and the minimal-copy baseline (both 53.6%). StoiAgent also achieves higher top-3 and top-5 accuracy than StoPred, with 85.7% (48/56) versus 80.4% (45/56) at top-3 and 87.5% (49/56) versus 85.7% (48/56) at top-5. These comparisons show that StoiAgent matches StoPred in top-1 accuracy while recovering more correct stoichiometries among its top-3 and top-5 ranked candidates.

**Figure 4:**
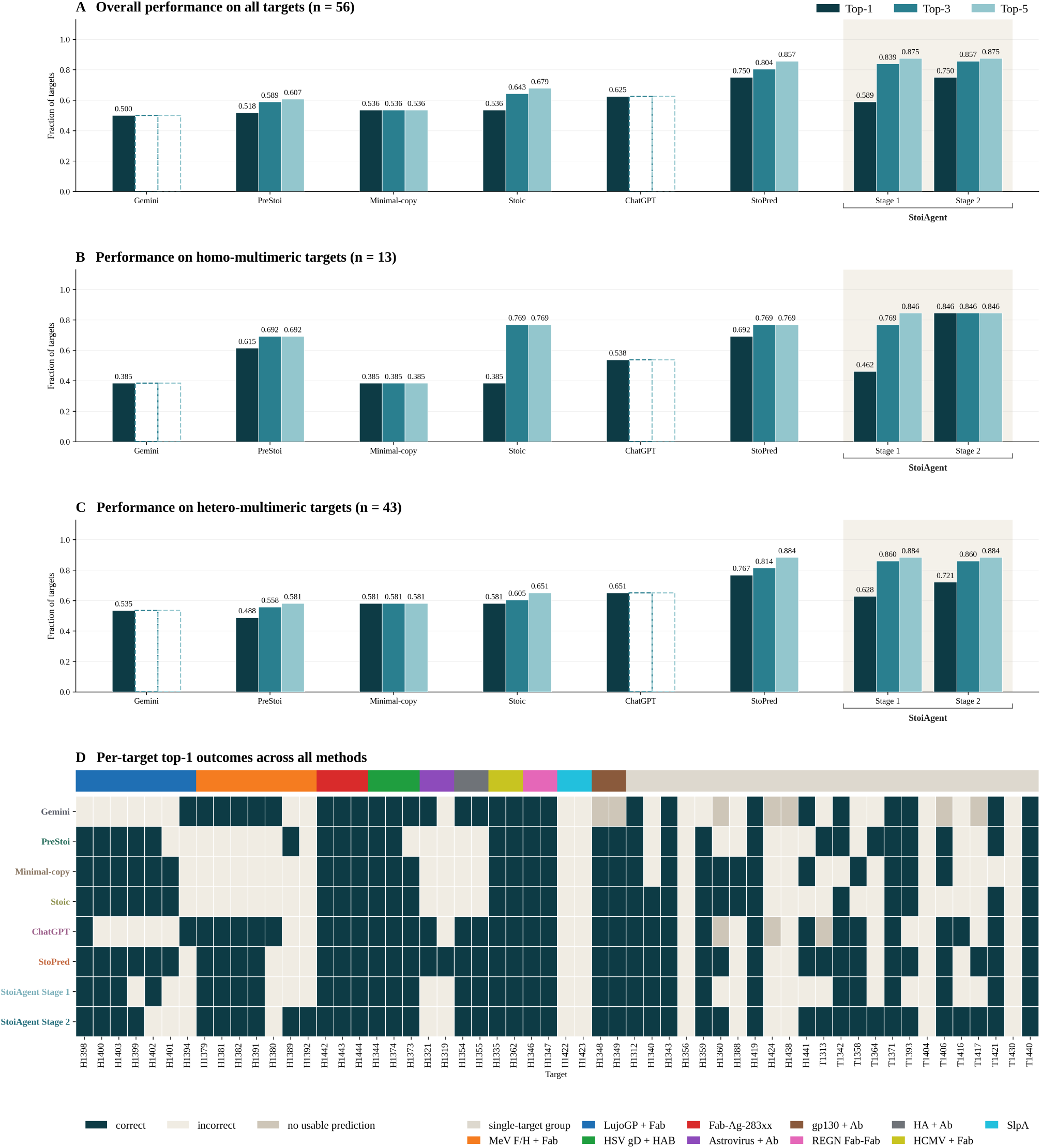
Comparison of protein complex stoichiometry prediction performance across methods. (A–C) Top-1 accuracy, top-3 accuracy, and top-5 accuracy for all 56 targets (A), 13 homomeric targets (B), and 43 heteromeric targets (C). Methods include the minimal-copy baseline, PreStoi based on AlphaFold3 ranking, StoiAgent in Stages 1 and 2, Stoic, StoPred, ChatGPT, and Gemini. ChatGPT and Gemini were evaluated using only protein sequences and target descriptions, and only top-1 accuracy is reported for them because they usually predict only one stoichiometry for a target. (D) Per-target top-1 prediction outcomes across all methods, grouped by biological system. Dark cells indicate correct predictions, light cells indicate incorrect predictions, and gray cells indicate unavailable or unusable predictions. All performance metrics use the full number of targets in the corresponding evaluation group as the denominator. For ChatGPT and Gemini, dashed bars represent visual placeholders at their Top-1 accuracy, as these models provided only single direct predictions.

To examine differences between complex types, we separately analyze homomeric and heteromeric targets (Fig. 4B,C). StoiAgent (Stage 2) achieved top-1 accuracies of 84.6% (11/13) for homomers and 72.1% (31/43) for heteromers. Compared with StoPred, it correctly predicts two additional homomeric targets (11 versus 9) but two fewer heteromeric targets (31 versus 33). At top-3, StoiAgent predicts 11/13 homomeric and 37/43 heteromeric native stoichiometries correctly, higher than 10/13 and 35/43 of StoPred, respectively. Thus, despite identical overall top-1 accuracy on all 56 targets, the two methods differ in their results of the two complex types. Per-target outcomes are shown in Fig. 4D.

We additionally evaluate ChatGPT and Gemini as general-purpose LLM baselines by providing the protein sequences and target descriptions to them and asking each model to predict the complex stoichiometry directly. ChatGPT achieved a top-1 accuracy of 62.5% (35/56), while Gemini achieved 50.0% (28/56), compared with 75.0% (42/56) for StoiAgent (Stage 2) (Fig. 4A). They also perform worse than StoiAgent (Stage 2) on either homomers or hteromers (Fig. 4B,C). The results demonstrate that the specialized StoiAgent based on a small open weight LLM outperforms the large frontier AI models through incorporating domain-specific inputs such as structural templates and AlphaFold3 ranking scores.

To account for the unequal representation of biological systems, we further calculate group-balanced metrics (Table 1). Across 34 systems, StoiAgent (Stage 2) achieves the highest group-balanced top-1 accuracy of 74.8%, compared with 51.5% for PreStoi with AlphaFold3 ranking, 71.8% for StoPred, 60.3% for ChatGPT, and 45.2% for Gemini. Its group-balanced top-3 and top-5 accuracy reaches 79.4% and 82.4%, respectively, exceeding StoPred’s 75.6% and 80.5%.

**Table 1:**
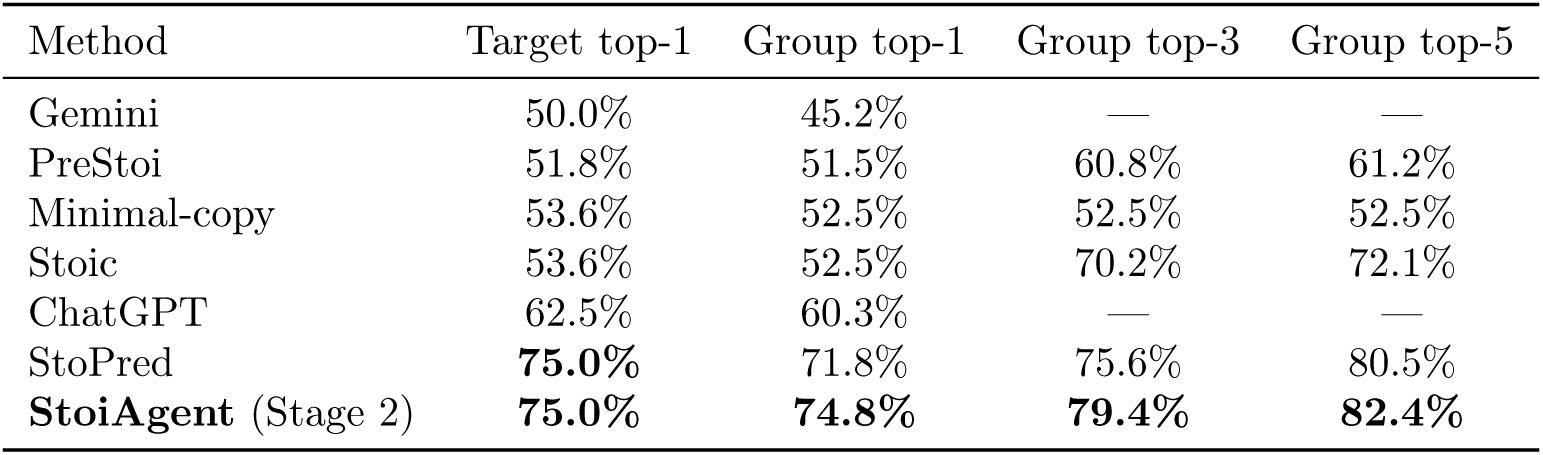
Target-level and group-balanced stoichiometry prediction performance for 56 CASP17 targets belonging to 34 annotated biological systems. Group-balanced metrics average within-system top-*k* accuracy and then assign equal weight to each system. Target-level top-1 accuracy is included for comparison with Fig. 4.

| Method | Target top-1 | Group top-1 | Group top-3 | Group top-5 |
|---|---|---|---|---|
| Gemini | 50.0% | 45.2% | — | — |
| PreStoi | 51.8% | 51.5% | 60.8% | 61.2% |
| Minimal-copy | 53.6% | 52.5% | 52.5% | 52.5% |
| Stoic | 53.6% | 52.5% | 70.2% | 72.1% |
| ChatGPT | 62.5% | 60.3% | — | — |
| StoPred | **75.0%** | 71.8% | 75.6% | 80.5% |
| **StoiAgent** (Stage 2) | **75.0%** | **74.8%** | **79.4%** | **82.4%** |

### StoiAgent effectively generates candidate stoichiometries in Stage 1

In Stage 1, StoiAgent places the native stoichiometry among its top-3 candidates for 47/56 CASP17 targets (83.9%) and among its top-5 for 49/56 targets (87.5%), while achieving a top-1 accuracy of 33/56 (58.9%). This difference indicates that StoiAgent (Stage 1) frequently includes the correct stoichiometry in top candidates but does not always rank it first.

To further assess candidate coverage, we examines the full Stage-1 candidate pool, which contains the native stoichiometry for 53/56 targets (94.6%). All 53 correct stoichiometries (references) appear within the top 10 positions, including three at rank 6 and one at rank 7. Thus, the fixed top-5 shortlist retains 49 of the 53 references generated by Stage 1 (92.5%). Among the 7 targets whose references are absent from the shortlist, four are excluded by the top-5 cutoff, while 3 are not generated at all. These results distinguish the limitations in candidate prioritization from those in candidate generation.

To illustrate how StoiAgent constructs alternative stoichiometry hypotheses in Stage 1, we examine the LujoGP + Fab9 complex (H1398, an antibody-antigen complex) as an example (Fig. 5). This target contains three unique protein subunits: Lujo viral glycoprotein A and antibody chains B and C. Several structural templates, including 8SAK, 9BTV, 8UZC, 8EEV, and 8TL4, showed significant sequence matches, with e-values ranging from 2.40 *×* 10*^−^*^17^ to 4.30 *×* 10*^−^*^15^. However, these templates cover only antibody chains B and C, leaving the copy number of glycoprotein A unresolved. The templates also exhibit different B:C stoichiometric patterns, including B1C1, B3C1, B3C3, and B6C3, providing incomplete and potentially conflicting evidence for the complete assembly.

**Figure 5:**
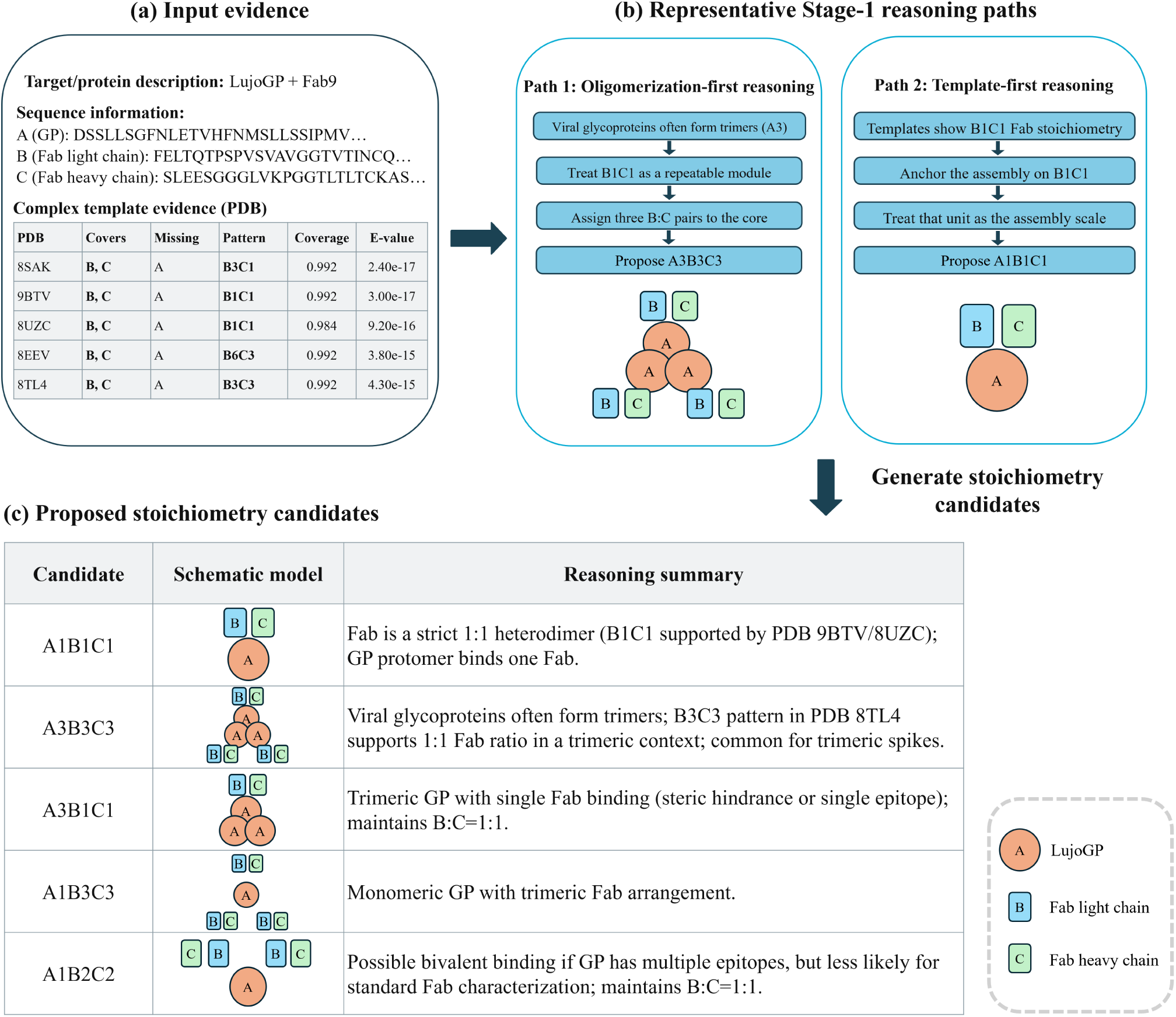
Evidence-guided candidate generation by StoiAgent for the LujoGP + Fab9 complex in Stage 1. (a) Input sequences and structural-template evidence, including mapped subunits, template stoichiometries, sequence coverage, and E-values. (b) Representative reasoning paths based on glycoprotein oligomerization and template-associated antibody-chain composition. (c) Five candidate stoichiometries with their schematic assemblies and supporting reasoning drawn from the Stage-1 trace. The alternative hypotheses reflect assumptions about the complete assembly that are not uniquely determined by the available templates.

The reported reasoning paths in Fig. 5 illustrate how different interpretations of the same evidence can produce alternative stoichiometries. One path assumes a trimeric glycoprotein associated with three B1C1 antibody pairs, leading to A3B3C3. Another prioritizes the B1C1 antibody module observed in structural templates and proposes A1B1C1. Additional candidates, including A1B2C2, A2B2C2, and A3B1C1, reflect different assumptions about glycoprotein oligomerization and antibody occupancy. Because the templates do not directly cover glycoprotein A, none of these arrangements is uniquely established by template evidence alone. This example illustrates how StoiAgent generates multiple candidate stoichiometries in Stage 1 when structural evidence is incomplete.

### StoiAgent improves prediction accuracy through stoichiometry reranking in Stage 2

In Stage 2, StoiAgent reranks the fixed top-five candidate shortlist using target information, Stage-1 ranks and support scores, and additional AlphaFold3 ranking scores (Fig. 6). Across the 56 CASP17 targets, top-1 accuracy increases from 33/56 (58.9%) in Stage 1 to 42/56 (75.0%) in Stage 2, a gain of 16.1 percentage points. StoiAgent corrects ten errors from Stage 1 while introducing one new error. Top-3 accuracy increases from 47/56 (83.9%) to 48/56 (85.7%), whereas top-5 accuracy remains at 49/56 (87.5%) because the five-candidate shortlist is unchanged. Thus, the improvement reflects more accurate selection among existing candidates by incorporating AlphaFold3 scores into candidate reranking.

**Figure 6:**
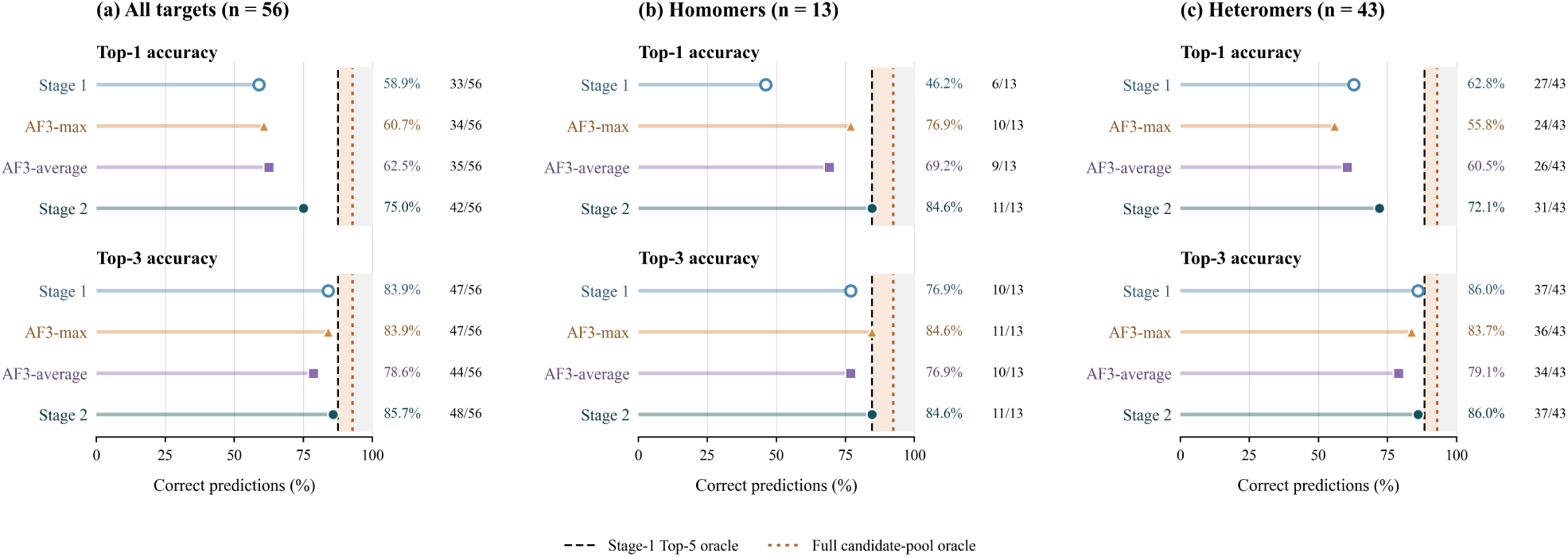
Comparison of stoichiometry reranking strategies on the fixed Stage-1 top-5 candidate shortlist. (a–c) Top-1 accuracy and top-3 accuracy for all 56 CASP17 targets (a), 13 homomeric targets (b), and 43 heteromeric targets (c). Four strategies are compared: the original Stage-1 ranking, ranking by maximum AlphaFold3 score (AF3-max), ranking by average AlphaFold3 score (AF3-average), and StoiAgent Stage 2. Black dashed lines indicate reference inclusion within the top-5 shortlist, while orange dashed lines indicate reference inclusion within the full Stage-1 candidate pool. These dotted lines represent theoretical accuracy ceilings based on candidate availability, rather than observed ranking accuracy. Numeric labels indicate correct-target counts.

To determine whether AlphaFold3 ranking scores alone could explain the improvement, we compare StoiAgent (Stage 2) with two baseline strategies that rank the same candidates using maximum or average AlphaFold3 scores (AF3-max and AF3-average). AF3-max and AF3-average achieve top-1 accuracies of 34/56 (60.7%) and 35/56 (62.5%), respectively, substantially lower than 42/56 (75.0%) of StoiAgent (Stage 2) (Fig. 6a). Their corresponding top-3 accuracy is 47/56 (83.9%) and 44/56 (78.6%), lower than 48/56 (85.7%) of StoiAgent (Stage 2). The results indicate that combining AlphaFold3 ranking scores with other complementary evidence by StoiAgent is important for accurate stoichiometry prediction.

The advantage of Stage 2 of StoiAgent is observed in both complex types. For heteromers, StoiAgent (Stage 2) achieves 31/43 (72.1%) top-1 accuracy, substantially higher than 24/43 (55.8%) for AF3-max and 26/43 (60.5%) for AF3-average (Fig. 6c). The same trent is observed for homomers (Fig. 6b). The comparisons indicate that StoiAgent (Stage 2) selects the native stoichiometry more accurately than the AlphaFold3 ranking score-based ranking strategies.

Despite the improved accuracy, the stoichiometry of 14 targets is not correctly predicted by StoiAgent in Stage 2. Seven errors result from correct stoichiometries being absent from the Stage-1 candidate shortlist, as described above. For the other seven, the correct stoichiometry is available but not selected as the top-ranked candidate; and all seven involve heteromeric complexes. For the 49 targets with their correct stoichiometry in the shortlist, StoiAgent achieves top-1 accuracy of 42/49 targets (85.7%) in Stage 2. These findings indicates there is still room of improvement in both stoichiometry candidate generation and ranking.

To illustrate stoichiometry reranking decisions, we examine two related LujoGP–Fab complexes with contrasting outcomes: H1398 (LujoGP + Fab9) and H1394 (LujoGP + Fab8) (Fig. 7). Their native stoichiometries are A1B1C1 and A3B3C3, respectively, even though their sequences are similar. The figure summarizes the key evidence and ranking considerations recorded in the Stage-2 reasoning traces.

**Figure 7:**
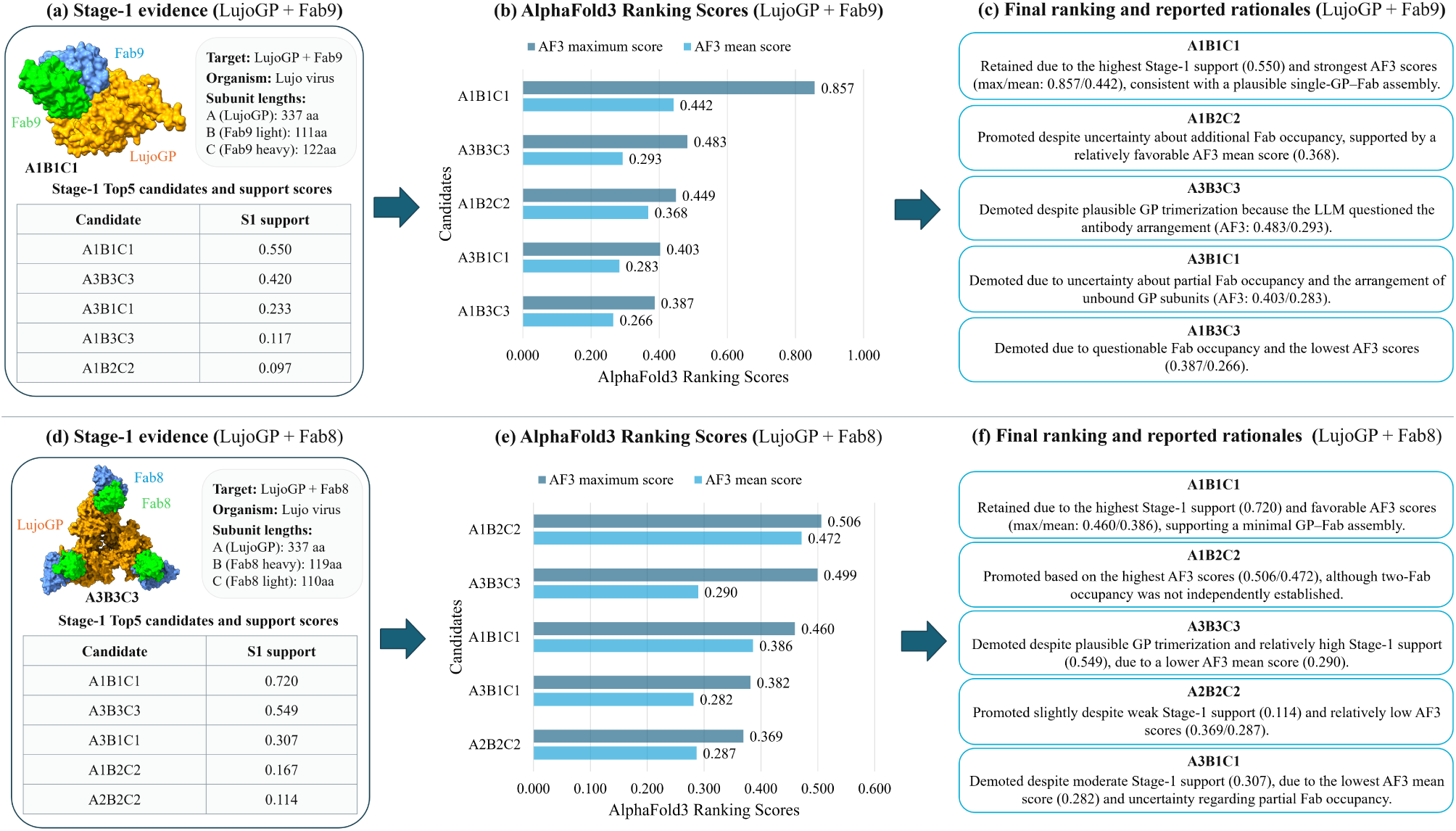
Contrasting Stage-2 reranking outcomes for two LujoGP–Fab complexes. (a–c) H1398 (LujoGP + Fab9), where the correct A1B1C1 stoichiometry remains ranked first in Stage 2. (d–f) H1394 (LujoGP + Fab8), where the reference A3B3C3 is demoted from second to third while the incorrect A1B1C1 remains first in Stage 2.

For H1398, the correct A1B1C1 composition is ranked first with a support score of 0.550 by StoiAgent in Stage 1. This candidate also has the highest AlphaFold3 maximum and mean scores (0.857 and 0.442), and StoiAgent retains it at rank 1 (Fig. 7a–c) in Stage 2. However, StoiAgent changes the ordering of alternative assemblies in Stage 2, promoting A1B2C2 from fifth to second and demoting A3B3C3 from second to third. The Stage-2 reasoning trace associates these changes with AlphaFold3 ranking scores and assumptions about antibody occupancy.

In contrast, H1394 illustrates a failure to select the correct stoichiometry despite its inclusion in the shortlist. StoiAgent ranks the incorrect A1B1C1 first with support 0.720, followed by the reference (correct) A3B3C3 with support 0.549 in Stage 1. Although A1B2C2 had the highest AlphaFold3 maximum and mean scores (0.506 and 0.472), in Stage 2, StoiAgent retains A1B1C1 at rank 1, promotes A1B2C2 from fourth to second, and demotes the reference A3B3C3 to third (Fig. 7d–f). The Stage-2 reasoning trace cites the relatively low mean AlphaFold3 score of A3B3C3 (0.290) as evidence against the trimeric assembly. Thus, the final ranking does not simply follow AlphaFold3 score ordering, but the additional considerations still does not produce the correct prediction.

To assess the effect of shortlist size, we further evaluate StoiAgent in Stage 2 using the top-10 Stage-1 candidates instead of the original top-5. Expanding the shortlist increases reference inclusion from 49/56 (87.5%) to 53/56 (94.6%). Final top-3 and top-5 accuracy increases from 48/56 (85.7%) and 49/56 (87.5%) to 50/56 (89.3%) and 51/56 (91.1%), respectively. However, top-1 accuracy decreases from 42/56 (75.0%) to 40/56 (71.4%). This comparison reveals a trade-off between reference coverage and final selection accuracy: expanding the candidate shortlist improves top-3 and top-5 accuracy but does not necessarily improve the top-1 accuracy because expanding the candidate shortlist makes it harder to select top-1 stoichiometry.

Overall, in Stage 2, StoiAgent improves candidate selection beyond simple AlphaFold3 score sorting, while remaining limited by shortlist coverage and some difficulty of distinguishing competing stoichiometries.

### Case studies of evidence-guided, interpretable stoichiometry prediction

To further investigate how StoiAgent integrates different sources of evidence, we examined two CASP17 targets: H1346 (REGN3753 Fab–Fab complex) and H1441 (NDM-1–Nanobody17 complex). These examples illustrate two distinct aspects of the workflow in StoiAgent: the influence of structural template evidence on candidate prioritization in Stage 1, and the ability of selecting the correct stoichiometry when both Stage-1 support and AlphaFold3 scores favor alternative ones in Stage 2.

For H1346, the native stoichiometry is A1B1C1D1, representing two Fab fragments, each comprising one heavy-chain fragment and one light chain. In a representative run, using only sequences and target descriptions, StoiAgent (Stage 1) incorrectly interpreted the Fab–Fab complex as requiring two copies of each supplied entity, ranking A2B2C2D2 first and the correct A1B1C1D1 second. This interpretation double-counted the constituent chains, as one copy of each entity already represents two Fab fragments. In contrast, when the structural template evidence was supplied, StoiAgent (Stage 1) ranked A1B1C1D1 first. The provided summaries for PDB entries 1FNS and 7KKZ supported this composition through mappings covering all four entities. It also identified the doubled A2B2C2D2 composition as an unsupported higher-order assembly. Because the correct stoichiometry was already present among the input candidates, the improvement was due to candidate prioritization rather than the generation of a new hypothesis. Notably, the template-supported run did not invoke external retrieval calls, indicating that the observed difference was associated with supplied template evidence rather than additional online research. The H1346 reasoning trace is provided in Supplementary Section 1.1 and 1.2.

H1441 illustrates a different scenario in which StoiAgent (Stage 2) correctly selected a stoichiometry that was not favored by either Stage-1 support or AlphaFold3 score sorting. StoiAgent (Stage 1) ranked A2B2 first with a support score of 0.787, while the reference A1B1 ranked second with a score of 0.500. All five shortlisted candidates had 100 AlphaFold3-predicted structures. A2B2 achieved the highest maximum AlphaFold3 ranking score (0.458), whereas A1B2 had the highest mean score (0.252). Despite its lower maximum and mean scores of 0.284 and 0.229, respectively, StoiAgent (Stage 2) promoted the correct A1B1 to the first place. The Stage-2 reasoning trace favored a one-to-one enzyme–nanobody binding arrangement and also noted that A1B1 had a higher mean AlphaFold3 score than A2B2 (0.229 versus 0.207), leading to the correct prediction. In contrast, neither retaining the original Stage-1 ranking nor sorting candidates by maximum or mean AlphaFold3 scores would have predicted the stoichiometry correctly. The H1441 trace is provided in Supplementary Section 1.3.

### Effects of LLM backbones on prediction accuracy

To assess the influence of LLM backbone selection on stoichiometry prediction, we evaluated four LLMs across all 56 CASP17 targets using the same Database + literature + templates configuration in Stage 1 and structural reranking procedure in Stage 2 (Fig. 8). The same as on CASP16 dataset, Qwen3.6-27B achieves the highest top-1 accuracy at 75.0% (42/56). It is followed by Gemma-3-27B-it at 69.6% (39/56), OLMo-2-32B-Instruct at 51.8% (29/56), and Mistral-Small-3.2-24B at 42.9% (24/56). Qwen maintains the highest top-1 and top5 accuracy, although its advantage is more pronounced in terms of top-1 accuracy.

**Figure 8:**
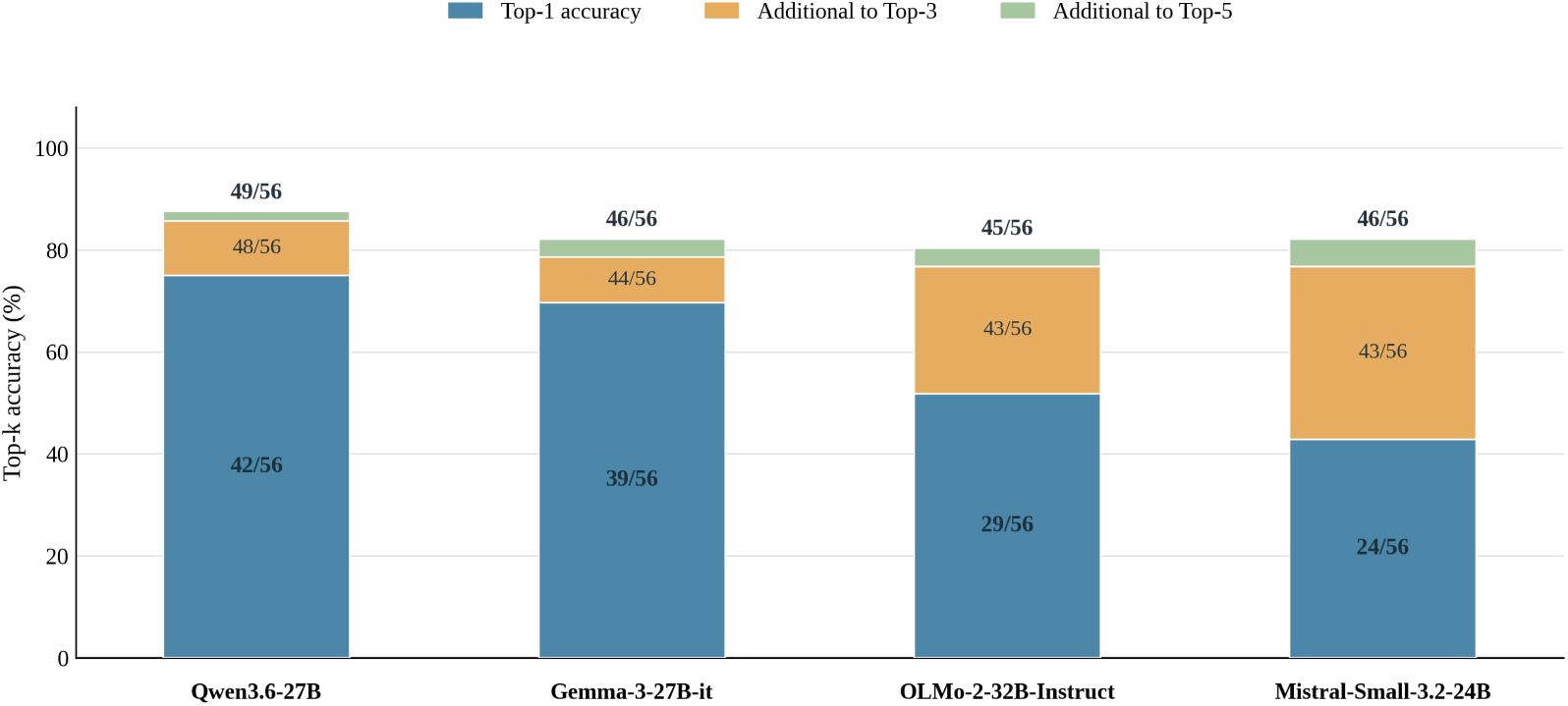
Comparison of Stage-2 stoichiometry prediction performance of StoiAgent with four different LLM backbones on 56 CASP17 targets. Qwen3.6-27B, Gemma-3-27B-it, OLMo-2-32B-Instruct, and Mistral-Small-3.2-24B were evaluated under the same Database + literature + templates configuration in Stage 1, followed by the reranking of the Stage-1 top-5 candidates using the AlphaFold3 ranking scores in Stage 2. Stacked bars show cumulative top-1, top-3, and top-5 accuracy. Blue segments show targets with a correct top-1 prediction; orange segments add targets whose correct stoichiometry is ranked second or third, and green segments add targets whose correct stoichiometry is ranked fourth or fifth. Numeric labels indicate cumulative correct-target counts out of 56. Missing or invalid predictions are counted as incorrect.

For Qwen and Gemma, the correct stoichiometry is ranked first for 42 of 49 and 39 of 46 targets, respectively, among those with the correct stoichiometry included in the top-5 shortlist. In contrast, the corresponding proportions are 29/45 for OLMo and 24/46 for Mistral. Qwen includes the correct stoichiometry in the top-5 shortlist for 49 targets, more than the other LLMs. Notably, Gemma and Mistral achieve the same top-5 accuracy (46/56, 82.1%), but their top-1 accuracy differs by 15 targets (39 versus 24). Similarly, OLMo and Mistral achieve comparable top-3 and top-5 accuracy despite differences in top-1 accuracy. These results indicate that the LLM backbones differ not only in their capability of including the correct stoichiometry in the top-5 shortlist but also in ranking it first.

## Discussion

StoiAgent uses an LLM-powered AI agent to integrate multiple sources of evidence including protein sequence, literature information, structural templates, and AlphaFold3 ranking scores to improve protein complex stoichiometry prediction. On the CASP17 benchmark, StoiAgent substantially outperforms PreStoi that similar structural template information and AlphaFold3 ranking scores but does not have the reasoning and integration provided by the LLM (top-1 accuracy: 75.0% versus 51.8%), indicating that the LLM reasoning is able to automatically integrate multi-sources of heterogeneous evidence to improve the prediction accuracy.

StoiAgent based on a small open-weight Qwen model also outperforms the two very large frontier AI models, ChatGPT (top-1 accuracy: 62.5%) and Gemini (top-1 accuracy: 50.0%) that are able to make stoichiometry prediction from only protein sequences and target descriptions. The results show that providing additional domain-specific information such structural templates and AlphaFold3 ranking scores can compensate the disadvantage of the small LLM to make it perform better than a large, advanced LLM.

Moreover, the overall prediction accuracy of StoiAgent without any training is better than StoPred, a deep learning model that was trained on a large set of protein complexes to predict stoichiometry from sequences alone. On one hand, the results indicate that integrating multiple sources of information such as structural templates, literature, and AlphaFold3 ranking scores is useful for improving stoichiometry prediction. On the other hand, the strong performance of StoPred shows that training AI models on protein complexes with known stoichiometry information is also important for stoichiometry prediction. Therefore, in the future, it is necessary to use known protein complexes to train AI agents such as StoiAgent to make it better in learning the rules of integrating heterogeneous to further improve the prediction accuracy.

A key feature of StoiAgent is its two-stage design, which separates the generation of plausible stoichiome-tries from their final prioritization. In Stage 1, it includes the native stoichiometry within the top-5 candidates for 49 out of 56 CASP17 targets (87.5%), but only correctly rank the native stoichiometry of 33 targets first. In Stage 2, it is able to substantially increase the number to 42 by incorporating AlphaFold3 ranking scores as additional evidence, demonstrating the structural modeling scores from AlphaFold3 is valuable for improving stoichiometry prediction. Moreover, StoiAgent (Stage 2) substan-tially outperforms the simple approach of ranking the same stoichiometry candidates by maximum or mean AlphaFold3 scores alone (top-1 accuracy: 75.0% versus 60.7 or 62.5%), clearly demonstrating the LLM reasoning helps integrate the AlphaFold3 scores with other evidence synergistically to improve stoichiometry prediction accuracy.

The effectiveness of reranking in Stage 2 also depends on the quality and size of the candidate shortlist. Expanding the Stage-2 shortlist from 5 to 10 candidates increases the native stoichiometry inclusion from 49/56 (87.5%) to 53/56 (94.6%) and also improves final top-3 and top-5 accuracy. However, top-1 accuracy decreases from 75.0% to 71.4%. This trade-off suggests that expanding the list of candidates has two conflicting effects: on one hand, it increases the chance of including the true stoichiometry into consideration; on the other hand, it makes it more challenging to rank the correct stoichiometry at the top because more candidates need to be considered.

The LLM backbone comparison shows that the effectiveness of evidence integration depends on LLMs. The selected Qwen3.6-27B model achieves the highest 75.0% Stage-2 top-1 accuracy, followed by Gemma-3-27B-it (69.6%), OLMo-2-32B-Instruct (51.8%), and Mistral-Small-3.2-24B (42.9%). Because each backbone generated its own Stage-1 candidate shortlist, these differences reflect the combined effects of candidate generation and Stage-2 reranking rather than an isolated measure of evidence integration.

One unique advantage of StoiAgent is that it records the trace of reasoning (chain of thoughts) in the process of integrating heterogeneous, sometime conflicting evidence to predict stoichiometry, which makes its prediction more interpretable than traditional deep learning methods. For instance, StoiAgent (Stage 2) correctly selects A1B1 for target H1441 despite higher Stage-1 support and maximum AlphaFold3 scores favoring A2B2. The Stage-2 reasoning trace shows that the higher mean structural score of A1B1 relative to A2B2 and biological considerations favoring a one-to-one enzyme–nanobody interaction are weighed by StoiAgent more than other factor to reach the correct prediction.

Despite the progress made by StoiAgent, there are several limitations that warrant consideration. First, the evaluation included only 56 targets representing 34 biological systems, with some systems contributing multiple targets. Larger and more diverse benchmarks, together with uncertainty estimates accounting for biological-system dependence, are needed to further assess generalizability in the future. Second, although the ablation experiments identify performance differences among input evidence configurations, the contributions of individual evidence sources to specific ranking decisions are not systematically isolated. Finally, computational cost remains a limitation, primarily due to AlphaFold3 structure predictions required to obtain ranking scores for each candidate stoichiometry. Future work should prioritize more effective candidate selection, efficient structural evaluation, improved integration of potentially conflicting evidence, and systematic assessment of evidence contributions.

## Methods

### StoiAgent workflow and two-stage stoichiometry prediction

StoiAgent is a two-stage LLM-powered framework for predicting protein complex stoichiometry (Fig. 1). Given the amino acid sequences of the unique subunits of a protein complex target and its simple description (e.g., protein name and organism information), the pipeline first gathers structural-template evidence relevant to those subunits. For heteromeric targets, templates matching multiple subunits are prioritized because they may show how the subunits assemble together. If such a template is unavailable, individual-subunit templates are used; these are also used for homomeric targets. The templates provide clues about possible copy numbers, but a partial match does not establish the stoichiometry of the entire target.

Specifically, in Stage 1, the LLM begins with the target information and template evidence to propose possible stoichiometries. It then retrieves supporting information from PDB, UniProt, and the literature, and reassesses its hypotheses in light of the retrieved evidence. This reasoning, retrieval, and synthesis cycle is represented by the connected steps in Fig. 1. The agent considers where evidence agrees or conflicts, and which subunit copy numbers remain uncertain. It produces a ranked list of complete stoichiometries, each with a support score. For example, A3B2C2 and A3B3C3 are two distinct hypotheses about the numbers of B and C subunits even though both contain three copies of A. The five highest-ranked candidates form the shortlist for Stage 2.

In Stage 2, AlphaFold3 is used to evaluate the structural plausibility of the shortlisted candidates by generating a number of structural models for each candidate and assessing the maximum or mean ranking scores of the models. The LLM integrates the resulting AlphaFold ranking scores with the Stage-1 support scores to rerank the candidates without introducing or removing any stoichiometries. Both stages preserve reasoning traces documenting the LLM’s chain of thoughts in making the ranking decisions. The specific evidence configurations and implementation details are described in the following subsections. The prompts used by StoiAgent in both stages are provided in Supplementary Section 2.

### Model development on CASP16 dataset and model evaluation on CASP17 dataset

We use 28 CASP16 Phase-0 targets, including 19 heteromeric and 9 homomeric complexes, to compare four LLM backbones and different evidence configurations. Based on Stage-2 top-1 accuracy, Qwen3.6-27B with the Database + literature + templates configuration is selected as the reasoning and action LLM for StoiAgent. The LLM is used as it is without being retrained.

StoiAgent with the selected setting is evaluated on CASP17 complex targets and compared with existing stoichiometry prediction methods. The evaluation is restricted to the 56 CASP17 multimer targets in the H and T series for which both an initial CASP17 Phase-1 target release without stoichiometry information and a corresponding CASP17 Phase-2 target release with stoichiometry information are available. Canceled targets are excluded. All predictions are evaluated against the true stoichiometries through exact matching, with missing or invalid predictions counted as incorrect. A prediction is considered correct only if the number of copies of every distinct protein subunit matches the reference one exactly.

It is worth noting that StoiAgent predictions were made after the official CASP17 experiment was concluded, even though it only used the information available prior to the release date of each CASP17 target, the evaluation is a retrospectively blind test instead of a prospectively blinded assessment.

### Target-specific temporal evidence restrictions

To simulate the blind prediction conditions of CASP17, we restricted external evidence retrieval to information available before each target’s initial release date. For each of the 56 CASP17 targets, the cutoff was defined as the day preceding its initial release. Since the targets were released between April 29 and July 17, 2026, the corresponding evidence cutoffs ranged from April 28 to July 16, 2026.

These restrictions were applied to the external databases accessed during Stage 1, including PDB, UniProt, and PubMed. PDB entries were accepted only if both their initial release and latest revision dates preceded the target-specific cutoff. UniProt records were filtered using their initial publication, sequence-update, and annotation-update dates. For PubMed, publication dates and record creation and revision dates were checked before the corresponding abstracts were made available to the LLM. Records with missing or unverifiable dates, or updates after the cutoff, were excluded. Structural template evidence was prepared separately using a PDB-derived database with a fixed cutoff of April 1, 2026. Stage 2 did not perform additional external retrieval.

These temporal restrictions were intended to prevent the agent from accessing information released after the CASP17 targets.

### Evidence configurations and Stage-1 stoichiometry candidate generation

In Stage 1, StoiAgent uses the target description and amino acid sequences of the unique protein subunits to generate and rank candidate stoichiometries. We evaluated two evidence configurations: Database + literature, and Database + literature + templates. Both evidence configurations use protein sequences and target descriptions as common inputs. The first configuration allows the LLM to retrieve biological evidence from PDB, UniProt, and PubMed. The second further provides structural-template evidence obtained through sequence-based template searches. During Stage 1, the LLM can selectively search the RCSB PDB by text or subunit sequence, search UniProt for protein annotations, and search PubMed for relevant literature. It can then inspect individual PDB and UniProt records or request PubMed records. The retrieved PubMed literature text is summarized by a separate, fixed Qwen3-4B-Instruct-2507 model. PDB and UniProt records and the literature summaries are returned to StoiAgent as tool observations.

Structural templates are identified using HH-suite 3.2.0 [22] against a PDB-derived template database [23]. For CASP17, this database was prepared using a cutoff date of April 1, 2026. HHmake is used to construct hidden Markov models from target sequence alignments, and HHsearch is used to identify homologous structural templates. Template hits are filtered according to sequence coverage, alignment length, and sequence similarity. Up to five highest-ranked templates are retained for each individual subunit in a target, while additional hits are examined to identify templates shared by multiple subunits.

For heteromeric complexes, templates matching at least two distinct subunits are prioritized because they can provide information about their assembly organization. When no eligible shared template is available, individual subunit templates are supplied to StoiAgent instead. Homomeric complexes receive individual subunit templates because there is only one unique subunit. Target-to-template chain correspondence is established through sequence matching, and subunit copy-number information is extracted from the corresponding PDB annotations of the templates. Template evidence is treated as supporting information rather than a definitive constraint on the target stoichiometry, particularly when a template covers only part of the complex.

For each target, in Stage 1, StoiAgent generates and ranks candidate stoichiometries in 15 independent runs with seeds 1234–1248, a temperature of 0.7, and top-*p* of 0.9. Each run is instructed to propose at least five distinct candidates, including the simplest stoichiometry (i.e., A2 for homomultimer or one copy for each unique subunit for heteromultimer). Incomplete or duplicated stoichiometries are removed. If a run generates less than five valid stoichiometries, StoiAgent is invoked to generate more until five is reached. The LLM-reported support scores for each stoichiometry candidate are averaged across the 15 runs, assigning zero to candidates absent from an individual run. Candidates are ranked by their average support scores. The five highest-ranked candidates form the fixed shortlist for Stage 2.

### AlphaFold3 structural evaluation and Stage-2 stoichiometry reranking

The complex structures of stoichiometry candidates in Stage 2 are predicted using AlphaFold3 version 3.0.2 (Git commit f6a5aec) [2]. For each candidate stoichiometry, 100 structures are generated using different random seeds. Candidates exceeding 5,000 total residues are excluded from AlphaFold3 structure prediction. For heteromultimers, the maximum and mean AlphaFold3 ranking score of the structures generated for each candidate are used as its structural scores. For homomultimers, only the maximum AlphaFold3 ranking score is used.

In Stage 2, StoiAgent considers the target information (sequence, name, and organism), the top-5 shortlist from Stage 1, the corresponding Stage-1 support scores, and the available AlphaFold3 ranking scores to rerank the stoichiometry candidates. Specifically, the LLM evaluates the competing stoichiometries using all the information and produces a revised ranking. Candidates without available AlphaFold3 scores remain in the shortlist and are treated as lacking structural evidence rather than being assigned an unfavorable score.

In Stage 2, StoiAgent records the reasoning traces that document the biological and structural evidence considered for the shortlisted candidates, the chain of thoughts, and the resulting ranking decisions made by the LLM.

### LLM backbones and baseline methods

Four LLM backbones were evaluated: Qwen3.6-27B, Gemma-3-27B-it, OLMo-2-32B-Instruct, and Mistral-Small-3.2-24B. The models were compared using identical prompts and evidence configurations. Qwen3.6-27B was selected as the primary backbone based on its better performance on the CASP16 dataset.

On the CASP17 test dataset, StoiAgent is compared with several existing stoichiometry prediction methods, including PreStoi, StoPred, Stoic, and the minimal-copy baseline. The stoichiometry predictions from PreStoi, Stoic and StoPred were made by using their open-source software released on GitHub.

ChatGPT and Gemini were also evaluated as general-purpose frontier LLM baselines. Both LLMs received protein sequences and target descriptions as input to predict the stoichiometry directly. Their predictions were recorded before the reference stoichiometries became available during the 2026 CASP17 experiment. Because ChatGPT and Gemini usually predicted only one stoichiometry for a target, only top-1 accuracy was evaluated.

## Data availability

The curated CASP16 and CASP17 benchmark datasets used in this study are publicly available at https://github.com/BioinfoMachineLearning/StoiAgent/tree/main/examples. The datasets con-tain 28 CASP16 and 56 CASP17 protein complex targets, including their amino acid sequences, target descriptions, and experimentally determined stoichiometries obtained from the official CASP releases. The repository also provides the original CASP data files and scripts used to construct the benchmark datasets.

## Code availability

The source code of StoiAgent is publicly available at https://github.com/BioinfoMachineLearning/ StoiAgent under the MIT license. The repository includes the implementation of the two-stage stoichiometry prediction framework, structural template search, AlphaFold 3 integration, and instructions for installation and execution. External databases and AlphaFold 3 model parameters must be obtained separately.

## Acknowledgements

We thank CASP organizers for making the CASP16 and CASP17 data available. This work is partially supported by an NIH NIGMS MIRA grant (#: R35GM161543).

## Author contributions

J.C. conceived the project. J.C., J.L., and P.N. designed the experiment. J.L., P.N., and J.C. performed the experiment and collected the data. J.L. and J.C. analyzed the data. J.L. and J.C. wrote the manuscript.

