## Supplementary document for "Accurate and interpretable protein complex stoichiometry prediction through agentic AI with StoiAgent"

November 17, 2026

### Supplementary Section 1. Representative StoiAgent Reasoning Traces

#### 1.1 H1346: REGN3753 Fab-Fab complex

**System prompt.** You predict the biological stoichiometry (copy number of each distinct protein subunit) of a protein complex. The distinct protein subunits are A, B, C, and D. Use only the target description/source and amino-acid sequences supplied by the user. Do not assume access to databases, templates, literature, or external tools.

**Stage-1 output policy.**

- Propose at least five complete and unique stoichiometry candidates.
- Include the simple baseline candidate `A1B1C1D1` so Stage 2 can compare it.
- Every candidate must contain every target subunit exactly once in canonical subunit order, with a positive integer copy number.
- For a single-subunit complex, `X1` is invalid.
- Give each candidate a support score from 0 to 1 based only on the evidence available in this Stage-1 condition.
- The Final Answer should be the best Stage-1 candidate; downstream Stage 2 will re-rank the candidate pool using AlphaFold3 ranking scores.

**Original target information supplied by the user.**

- Target/protein description: REGN3753 Fab-Fab complex
- Organism/source: Homo sapiens
- Distinct protein subunits:
  - Subunit A: released sequence record/subunit 1; length = 226 residues
    `QVQLQQWGAGLLKPSETLSLTCAVYGGSLSDYYWTWIRQPPEKGLEWIGEINHSGSTNYAPSLKSRVTI`
    `SVDTSKNQFSLKLTSVTAADTAVYYCARTTYGYDYYFDMDVWGQGTTVTVSSAKTTAPSVYPLAPVCGD`
    `TTGSSVTLGCLVKGYFPEPVTLTWNSGSLSSGVHTFPAVLQSDLYTLSSSVTVTSSTWPSQSITCNVAH`
    `PASSTKVDKKIEPRGPTIK`
  - Subunit B: released sequence record/subunit 2; length = 214 residues
    `DIQMTQSPSSLSASVGDRVTITCRASQVITNFLAWYQQTPGKVPKLLIYAASTLQSGVPSRFSGSGSGT`
    `DFTLTISSLQPEDVATYYCQKYNYTPLTFGGGTKVEIKRADAAPTVSIFPPSSEQLTSGGASVVCFLNN`
    `FYPKDINVKWKIDGSERQNGVLNSWTDQDSKDSTYSMSSTLTLTKDEYERHNSYTCEATHKTSTSPIVK`
    `SFNRGEC`

- Subunit C: released sequence record/subunit 3; length = 230 residues
  EVQLVESGGGLVQPGGSLRLSCAASGFTFSNFWMTWVRQAPGKGLEWVANMNQDGSEKYYVDSVKGRFT
  ISRDNAKSSLYLQMNSLRAEDTAVYYCARDREYCISTSCYDDFDYWGQGTLVTVSSASTKGPSVFPLAP
  CSRSTSESTAALGCLVKDYFPEPVTVSWNSGALTSGVHTFPAVLQSSGLYSLSSVVTVPSSSLGTKTYT
  CNVDHKPSNTKVDKRVESKYGPP
- Subunit D: released sequence record/subunit 4; length = 215 residues
  DIQMTQSPSSLSASVGDRVTITCRASQSISSYLNWYQQKPGKAPKLLIYAASSLQSGVPSRFSGSGSGT
  DFTLTISSLQPEDFATYYCQQSYSTPPITFGQGTRLEIKRTVAAPSVFIFPPSDEQLKSGTASVVCLLN
  NFYPREAKVQWKVDNALQSGNSQESVTEQDSKDSTYSLSSTLTLSKADYEKHKVYACEVTHQGLSSPVT
  KSFNRGEC

**User request.** Generate the Stage-1 candidate stoichiometries for downstream AlphaFold3 re-ranking.

**Assistant response.**

> **Thought:** The target is described as a "Fab-Fab complex," which structurally consists of two Fab fragments. Each Fab fragment is composed of one heavy chain variable/constant domain pair (Subunits A and C) and one light chain variable/constant domain pair (Subunits B and D). Therefore, the biological assembly requires two copies of each distinct subunit to form the two identical or heterologous Fab arms. The simplest stoichiometry A2B2C2D2 represents this dimeric Fab-Fab structure. Other candidates are proposed to cover potential oligomeric states or modeling artifacts, but A2B2C2D2 is the most biologically consistent with the "Fab-Fab" description.
>
> **Candidates:**
>
> 1. A2B2C2D2 support=0.95 support_evidence=Standard Fab-Fab dimeric structure implies two copies of each chain
> 2. A1B1C1D1 support=0.30 support_evidence=Baseline monomeric unit, though biologically incomplete for a Fab-Fab complex
> 3. A4B4C4D4 support=0.10 support_evidence=Potential higher-order oligomerization or aggregation artifact
> 4. A2B1C2D1 support=0.05 support_evidence=Unlikely asymmetric stoichiometry for a standard Fab-Fab complex
> 5. A1B2C1D2 support=0.05 support_evidence=Unlikely asymmetric stoichiometry for a standard Fab-Fab complex
>
> **Final Answer:** A2B2C2D2
>
> **Support Score:** 0.95

### 1.2 H1346: REGN3753 Fab–Fab complex with shared-template evidence

**System prompt.** You are an AI research agent that predicts protein-complex stoichiometry. Target subunits: A, B, C, D. Reason from the original target sequence/description, advisory shared/multi-subunit structural-template evidence, and selectively requested RCSB/UniProt/PubMed evidence. No isolated per-subunit template-copy evidence is provided in this condition. Shared-template evidence is not a hard constraint and must not override stronger target-specific evidence merely because it is present. Do not double-count multiple fields from the same PDB as independent evidence. Follow the common Stage-1 output format and never invent Observation lines.

**User request.** Predict the biological stoichiometry of this protein complex.

**Original target information.**

- Target/protein description: REGN3753 Fab-Fab complex
- Organism/source: Homo sapiens
- Subunit sequences:
  - Subunit A: CASP subunit 1; length = 226 residues
    QVQLQQWGAGLLKPSETLSLTCAVYGGSLSDYYWTWIRQPPEKGLEWIGEINHSGSTNYAPSLKSRVTI
    SVDTSKNQFSLKLTSVTAADTAVYYCARTTYGYDYYFDMDVWGQGTTVTVSSAKTTAPSVYPLAPVCGD
    TTGSSVTLGCLVKGYFPEPVTLTWNSGSLSSGVHTFPAVLQSDLYTLSSSVTVTSSTWPSQSITCNVAH
    PASSTKVDKKIEPRGPTIK
  - Subunit B: CASP subunit 2; length = 214 residues
    DIQMTQSPSSLSASVGDRVTITCRASQVITNFLAWYQQTPGKVPKLLIYAASTLQSGVPSRFSGSGSGT
    DFTLTISSLQPEDVATYYCQKYNYTPLTFGGGTKVEIKRADAAPTVSIFPPSSEQLTSGGASVVCFLNN
    FYPKDINVKWKIDGSERQNGVLNSWTDQDSKDSTYSMSSTLTLTKDEYERHNSYTCEATHKTSTSPIVK
    SFNRGEC
  - Subunit C: CASP subunit 3; length = 230 residues
    EVQLVESGGGLVQPGGSLRLSCAASGFTFSNFWMTWVRQAPGKGLEWVANMNQDGSEKYYVDSVKGRFT
    ISRDNAKSSLYLQMNSLRAEDTAVYYCARDREYCISTSCYDDFDYWGQGTLVTVSSASTKGPSVFPLAP
    CSRSTSESTAALGCLVKDYFPEPVTVSWNSGALTSGVHTFPAVLQSSGLYSLSSVVTVPSSSLGTKTYT
    CNVDHKPSNTKVDKRVESKYGPP
  - Subunit D: CASP subunit 4; length = 215 residues
    DIQMTQSPSSLSASVGDRVTITCRASQSISSYLNWYQQKPGKAPKLLIYAASSLQSGVPSRFSGSGSGT
    DFTLTISSLQPEDFATYYCQQSYSTPPITFGQGTRLEIKRTVAAPSVFIFPPSDEQLKSGTASVVCLLN
    NFYPREAKVQWKVDNALQSGNSQESVTEQDSKDSTYSLSSTLTLSKADYEKHKVYACEVTHQGLSSPVT
    KSFNRGEC

**Shared/multi-subunit structural-template evidence (advisory, not constraints).**

- Only templates that jointly map at least two target subunits are shown.
- Each listed PDB is one structural evidence source; fields derived from the same PDB are not independent evidence.
- A mapped biological assembly from a homologous/shared template may support a relative copy-number pattern, but it is not a hard constraint on the target.
- Do not assume that a partial shared-template pattern determines the copy number of uncovered target subunits.
- Target-specific external database/literature evidence may confirm or contradict shared-template evidence.

- PDB=1FNS; covers = [A, B, C, D]; observed pattern = [A1B1C1D1]; best coverage = 1.000; best E-value = 7.60e-31; type = FULL_COMPLEX_SHARED
- PDB=7KKZ; covers = [A, B, C, D]; observed pattern = [A1B1C1D1]; best coverage = 1.000; best E-value = 7.00e-30; type = FULL_COMPLEX_SHARED
- PDB=9XOU; covers = [A, B, C, D]; best coverage = 0.942; best E-value = 3.50e-28; type = FULL_COMPLEX_SHARED
- PDB=4UOM; covers = [A, B, C, D]; best coverage = 0.940; best E-value = 1.30e-33; type = FULL_COMPLEX_SHARED
- PDB=6W4V; covers = [A, B, D]; missing = [C]; observed pattern = [A1B1D1]; best coverage = 1.000; best E-value = 2.10e-28; type = MAJOR_PARTIAL_SHARED

**Optional external research.** Use external research selectively when it can resolve biological assembly, oligomerization, conflicting evidence, or target-specific identity. Available actions:

- `RCSB.search("text query")`
- `RCSB.searchSequence("A")`
- `RCSB.getPDB("1ABC")`
- `UniProt.search("text query")`
- `UniProt.get("P12345")`
- `PubMed.search("text query")`
- `PubMed.readAbstract("PMID")`

Do not invent Observation lines.

#### Stage-1 output policy.

- Propose at least five complete and unique stoichiometry candidates.
- Include the simple baseline candidate A1B1C1D1 so Stage 2 can compare the same baseline across variants.
- Every candidate must contain every target subunit exactly once in canonical subunit order, with a positive integer copy number.
- For a single-subunit complex, X1 is invalid.
- Give each candidate a support score from 0 to 1 based on the evidence available in this condition.
- Shared-template patterns are suggestions only; include or rank them only when judged biologically plausible.
- The Final Answer should be the best Stage-1 candidate; downstream Stage 2 will re-rank the candidate pool using AlphaFold3 ranking scores.

**Required final format.**

```
Thought: <1-3 short sentences>
Candidates:
1) <complete stoichiometry>  support=0.70  support_evidence=<short phrase>
2) <complete stoichiometry>  support=0.25  support_evidence=<short phrase>
IMPORTANT: support must be a bare numeric value from 0 to 1 with NO angle brackets.
Correct: support=0.70    Incorrect: support=<0.70>
3) ...
4) ...
5) ...
Final Answer: <one of the candidates>
Support Score: <support score of the Final Answer>
```

**Assistant response.**

**Candidates:**

1. `A1B1C1D1` support=0.950000 support_evidence=Consistent with Fab-Fab dimer structure (2 HC + 2 LC) and all shared templates
2. `A2B2C2D2` support=0.050000 support_evidence=Unlikely dimer of dimers without specific cross-linking evidence
3. `A1B1C1D2` support=0.010000 support_evidence=Asymmetric composition not supported by Fab symmetry
4. `A2B1C1D1` support=0.010000 support_evidence=Asymmetric composition not supported by Fab symmetry
5. `A1B2C1D1` support=0.010000 support_evidence=Asymmetric composition not supported by Fab symmetry

**Final Answer:** `A1B1C1D1`

**Support Score:** 0.950000

### 1.3 H1441: NDM-1–Nanobody17 stoichiometry reranking trace

**System prompt.** You are a scientific assistant reranking existing protein-complex stoichiometry hypotheses. Your job is to order the supplied candidate stoichiometries, not generate hypotheses.

**Hard constraints.**

- Return every supplied candidate exactly once, without additions, omissions, or changes.
- List the candidates in plain text, one stoichiometry per line, from most to least plausible.
- Put `Final Answer: <stoichiometry>` immediately after the candidate list. The Final Answer must equal the first-ranked candidate.
- A `Support Evidence` section may follow the ranking.
- Do not return JSON, Markdown fences, new candidates, or additional numbered lists.
- Do not browse or request external evidence, or assume access to evidence not shown.
- Write the complete ranked list and Final Answer before any extended explanation.

**Scientific interpretation.** Weigh the supplied target description, sequences, and Stage-1 support information. Stage-1 support is a heuristic, not a calibrated probability. Do not automatically select the Stage-1 first candidate or the simplest stoichiometry. The only valid output is a permutation of the input candidates; consider all of them.

**AlphaFold3 evidence rules.** AlphaFold3 `max_ranking_score` and `avg_ranking_score` measure predicted-structure confidence, not the probability that a biological stoichiometry is correct. Compare both scores. A high maximum with a much lower average may indicate variable confidence among generated models. Consider `num_models` when comparing scores; results from different numbers of predictions may not be directly comparable. Missing scores mean structural evidence is unavailable, not that the score is zero or that the candidate has negative biological evidence.

**Assembly-size interpretation rules.**

- Compare scores cautiously when candidates differ in total residue count, chain count, or copy number. A larger assembly introduces additional relative chain arrangements and potential interfaces, which can reduce global confidence even when the larger stoichiometry is biologically correct.
- A higher score for a smaller complex does not by itself establish that the smaller stoichiometry is biologically correct. Likewise, do not automatically prefer a larger candidate because it has more copies, greater symmetry, or more apparently occupied binding sites.
- Look for target-specific biological evidence supporting or contradicting the additional copies and interactions required by a larger candidate. If only global AlphaFold3 scores are supplied, do not infer interface scores, contacts, clashes, or other unprovided structural observations.
- Do not apply a fixed correction, bonus, penalty, or numerical normalization based on sequence length, total residues, chain count, or copy number. Assess size-related uncertainty when interpreting the raw scores.
- Neither the highest AlphaFold3 score nor the highest Stage-1 support must determine the ranking. If structural scores and biological evidence disagree, explain the tradeoff and uncertainty.

**User request.** Rerank the supplied candidate stoichiometries using the target information and evidence below. Return every candidate exactly once, then the Final Answer, followed by concise Support Evidence.

**Original target information.**

- Target/protein description: NDM-1 (ID C7C422): Nanobody17, *Klebsiella pneumoniae*
- Organism/source: *Camelus dromedarius*
- Subunit sequences:
  - Subunit A; length = 243 residues
    `MGEIRPTIGQQMETGDQRFGDLVFRQLAPNVWQHTSYLDMPGFGAVASNGLIVRDGGRVLVVDTAWTDD`
    `QTAQILNWIKQEINLPVALAVVTHAHQDKMGGMDALHAAGIATYANALSNQLAPQEGMVAAQHSLTFAA`
    `NGWVEPATAPNFGPLKVFYPGPGHTSDNITVGIDGTDIAFGGCLIKDSKAKSLGNLGDADTEHYAASAR`
    `AFGAAFPKASMIVMSHSAPDSRAAITHTARMADKLR`

– Subunit B; length = 134 residues
QVQLQESGGGSVQAGGSLRLSCKVSGNAASIGYLAWFRQAPGKEREGVAALYTGTGGTNYVDSLKGRFR
ISRDLPKNTLYLQMNNLKPEDSAMYYCARGMGMLFSVAEIFSPTNYNYWGQGTQVTVSSHHHHHH

**Stage-1 candidate pool.** Only the following candidates are allowed. Each candidate was evaluated using 100 AlphaFold3 models.

| Candidate | Stage-1 rank | Stage-1 support | AF3 max | AF3 average |
|---|---|---|---|---|
| A2B2 | 1 | 0.787 | 0.457738 | 0.206994 |
| A1B1 | 2 | 0.500 | 0.284481 | 0.228914 |
| A2B1 | 3 | 0.477 | 0.213681 | 0.193079 |
| A1B2 | 4 | 0.078 | 0.305963 | 0.251804 |
| A2B4 | 5 | 0.063 | 0.203058 | 0.180391 |

**Required output format.**

```
Candidates:
1. <stoichiometry>
2. <stoichiometry>
3. <stoichiometry>
4. <stoichiometry>
5. <stoichiometry>
Final Answer: <stoichiometry>
Support Evidence: <concise explanation>
```

**Assistant response.**

**Candidates:**

1. A1B1
2. A2B2
3. A1B2
4. A2B1
5. A2B4

**Final Answer:** A1B1

**Support Evidence:** The target describes a complex between NDM-1, a beta-lactamase enzyme, and Nanobody17. Nanobodies are single-domain antibodies derived from camelid heavy-chain-only antibodies, which typically bind to their antigen in a 1:1 stoichiometry. While NDM-1 is known to form dimers in solution, the specific interaction with a nanobody is most commonly characterized as a monomeric binding event per active site, or the nanobody stabilizes the monomeric form for structural determination.

Analyzing the provided data:

1. **Biological plausibility:** A1B1 is the standard stoichiometry for antibody–antigen complexes unless specific dimerization of the antibody or multivalent binding is indicated. Nanobodies are monovalent.

2. **AlphaFold3 scores:** A1B1 has a higher average ranking score (0.228914) than A2B2 (0.206994). Although A2B2 has a higher maximum score (0.457738 versus 0.284481), its lower average score suggests that predictions for the larger complex are less consistent or confident on average. The higher average score for A1B1 indicates a more robust prediction for the 1:1 complex.
3. **Stage-1 support:** Stage 1 favors A2B2 (0.787) over A1B1 (0.500), but the instructions state that Stage-1 support is a heuristic, not a calibrated probability. Given the biological precedent for 1:1 nanobody binding and the higher average AlphaFold3 confidence for A1B1, it is ranked first.
4. **Other candidates:** A1B2 and A2B1 are less likely given the monovalent nature of nanobodies and the typical dimeric nature of NDM-1. A2B1 is possible, but less likely than A1B1 for a specific nanobody interaction unless the nanobody bridges dimers, which is rare. A2B4 is highly unlikely given its low Stage-1 support and low AlphaFold3 scores.

Therefore, A1B1 is the most plausible stoichiometry, followed by A2B2, which might represent the dimeric enzyme with two bound nanobodies, but has lower average structural confidence.

### Supplementary Section 2. StoiAgent prompt templates

The following templates document the initial messages supplied to the LLM. Double-braced fields represent target-specific values or repeated records inserted by the scripts. The original target-information block contains the description, organism when available, and sequences of the unique subunits; it does not include the benchmark target identifier. Tool observations and validation-repair messages vary by run and are preserved in the corresponding execution traces.

#### 2.1 Stage 1: candidate generation

Stage 1 uses the shared-template prompt when a template maps at least two distinct target subunits. Otherwise, it uses the individual-subunit-template fallback prompt, including for homomeric targets.

**Target-information block used in both branches.**

```
0. Original target information:
- Target/protein description: {{DESCRIPTION}}
- Organism/source: {{ORGANISM}}
- Subunit sequences:
  - Subunit {{SUBUNIT_ID}}: {{SOURCE_SEQUENCE_LABEL}}; length={{LENGTH}} residues
    Sequence: {{AMINO_ACID_SEQUENCE}}
{{REPEAT_FOR_EACH_UNIQUE_SUBUNIT}}
```

##### 2.1.1 Shared-complex-template branch

**System message.**

```
You are an AI research agent that predicts protein-complex stoichiometry.
Target subunits: {{COMMA-SEPARATED_SUBUNIT_IDS}}.
Reason from the original target sequence/description, advisory shared/multi-subunit
    structural-template evidence, and selectively requested RCSB/UniProt/PubMed evidence.
```

```
No isolated per-subunit template-copy evidence is provided in this condition. Shared-template evidence is not a hard constraint and must not override stronger target-specific evidence merely because it is present.
Do not double-count multiple fields from the same PDB as independent evidence.
Follow the common Stage-1 output format and never invent Observation lines.
```

#### User message.

```
Predict the biological stoichiometry of this protein complex.

{{ORIGINAL_TARGET_INFORMATION_BLOCK}}

1. Shared/multi-subunit structural-template evidence (ADVISORY, not constraints):
- Only templates that jointly map at least two target subunits are shown here.
- Each listed PDB is one structural evidence source; fields derived from the same PDB are not independent evidence.
- A mapped biological assembly from a homologous/shared template may support a relative copy-number pattern, but it is not a hard constraint on the target.
- Do not assume that a partial shared-template pattern determines the copy number of uncovered target subunits.
- Target-specific external database/literature evidence may confirm or contradict shared-template evidence.
{{UP_TO_FIVE_SHARED_TEMPLATE_RECORDS}}

2. Optional external research:
- Use external research selectively when it can resolve biological assembly, oligomerization, conflicting evidence, or target-specific identity.
- Available Actions:
  Action: RCSB.search("text query")
  Action: RCSB.searchSequence("A")
  Action: RCSB.getPDB("1ABC")
  Action: UniProt.search("text query")
  Action: UniProt.get("P12345")
  Action: PubMed.search("text query")
  Action: PubMed.readAbstract("PMID")
- Do not invent Observation lines.

Stage-1 output policy:
- Propose at least 5 COMPLETE and UNIQUE stoichiometry candidates.
- Include the simple baseline candidate {{BASELINE_STOICHIOMETRY}} so Stage 2 can compare the same baseline across variants.
- Every candidate must contain every target subunit exactly once in canonical subunit order, with a positive integer copy number.
- For a single-subunit complex, X1 is invalid.
- Give each candidate a support score from 0 to 1 based on the evidence available in this condition.
- Shared-template patterns are suggestions only; include or rank them only when you judge them biologically plausible.
- The Final Answer should be your best Stage-1 candidate; downstream Stage 2 will re-rank the candidate pool using AlphaFold3 ranking scores.

Required final format:
Thought: <1-3 short sentences>
Candidates:
1) <complete stoichiometry>  support=0.70  support_evidence=<short phrase>
2) <complete stoichiometry>  support=0.25  support_evidence=<short phrase>
IMPORTANT: support must be a bare numeric value from 0 to 1 with NO angle brackets.
Correct: support=0.70    Incorrect: support=<0.70>
3) ...
4) ...
```

```
5) ...
Final Answer: <one of the candidates>
Support Score: <support score of the Final Answer>
```

The values 0.70 and 0.25 above are literal output-format examples in the script, not target-specific support scores.

#### 2.1.2 Individual-subunit-template fallback

**System message.**

```
You are an AI research agent that predicts protein-complex stoichiometry.
Target subunits: {{COMMA-SEPARATED_SUBUNIT_IDS}}.
Reason from the original target sequence/description, compact advisory structural-template
    evidence, and selectively requested RCSB/UniProt/PubMed evidence.
Template evidence is not a hard constraint and must not override stronger target-specific
    evidence merely because it is present.
Do not double-count derived summaries and their source template records as independent
    evidence.
Follow the common Stage-1 output format and never invent Observation lines.
```

**User message.**

```
Predict the biological stoichiometry of this protein complex.

{{ORIGINAL_TARGET_INFORMATION_BLOCK}}

1. Compact structural-template evidence (ADVISORY, not constraints):
- Template evidence is homolog/structure evidence and may not transfer exactly to the
    target biological assembly.
- Do NOT combine copy numbers from unrelated per-subunit templates into a full-complex
    stoichiometry automatically.
- Repeated summaries or multiple records from the same PDB are not independent evidence.
- A shared template covering multiple target subunits is usually more directly informative
    than an isolated subunit homolog, but neither is mandatory.
- Target-specific external database/literature evidence, when retrieved, may confirm or
    contradict template evidence.
{{PER_SUBUNIT_COPY_OBSERVATIONS_AND_REPRESENTATIVE_TEMPLATE_HITS}}

2. Shared/multi-subunit template evidence (ADVISORY):
- no shared multi-subunit template evidence

3. Optional external research:
- Use external research selectively when it can resolve biological assembly,
    oligomerization, conflicting evidence, or target-specific identity.
- Available Actions:
  Action: RCSB.search("text query")
  Action: RCSB.searchSequence("A")
  Action: RCSB.getPDB("1ABC")
  Action: UniProt.search("text query")
  Action: UniProt.get("P12345")
  Action: PubMed.search("text query")
  Action: PubMed.readAbstract("PMID")
- Do not invent Observation lines.

Stage-1 output policy:
- Propose at least 5 COMPLETE and UNIQUE stoichiometry candidates.
- Include the simple baseline candidate {{BASELINE_STOICHIOMETRY}} so Stage 2 can compare
    the same baseline across variants.
```

```
- Every candidate must contain every target subunit exactly once in canonical subunit order
    , with a positive integer copy number.
- For a single-subunit complex, X1 is invalid.
- Give each candidate a support score from 0 to 1 based on the evidence available in this
    condition.
- Template-derived patterns, scaffold completions, and symmetry patterns are suggestions
    only; include them only if you judge them biologically plausible.
- The Final Answer should be your best Stage-1 candidate; downstream Stage 2 will re-rank
    the candidate pool using AlphaFold3 ranking scores.

Required final format:
Thought: <1-3 short sentences>
Candidates:
1) <complete stoichiometry>  support=<0..1>  support_evidence=<short phrase>
2) <complete stoichiometry>  support=<0..1>  support_evidence=<short phrase>
3) ...
4) ...
5) ...
Final Answer: <one of the candidates>
Support Score: <support score of the Final Answer>
```

For each subunit, the variable template block reports the distribution of copy numbers among significant template hits and up to three representative hits. These are two presentations of the same underlying template evidence, not independent observations.

### 2.2 Stage 2: fixed-candidate reranking

The Stage-2 system message consists of the common instructions below, followed by the evidence rules appropriate to the target and condition. The user message then supplies the target and candidate-specific values.

**Common system message.**

```
You are a scientific assistant reranking existing protein-complex stoichiometry hypotheses.
Your job is to ORDER the supplied candidate stoichiometries, NOT generate hypotheses.

HARD CONSTRAINTS:
1. Return EVERY supplied candidate exactly once, without additions, omissions, or changes.
2. Return a plain-text Candidates section: one numbered stoichiometry per line, from most
    to least plausible.
3. Return Final Answer: <stoichiometry> immediately after the candidate list.
4. Final Answer MUST equal the first-ranked candidate.
5. You may then write a Support Evidence section explaining the decision.
6. Do not return JSON, Markdown fences, new candidates, or additional numbered lists.
7. Do not browse or request external evidence. Do not assume access to evidence not shown.
8. Write the complete ranked list and Final Answer BEFORE any extended explanation.

SCIENTIFIC INTERPRETATION:
- Weigh the supplied target description, sequences, and Stage-1 support information.
- The Stage-1 confidence/support is a heuristic, NOT a calibrated probability.
- Do not automatically select the Stage-1 first candidate or the simplest stoichiometry.
- The only valid output is a permutation of the input candidates; consider all of them.
```

**AlphaFold3 rules for heteromeric targets.**

```
ALPHAFOLD3 EVIDENCE RULES:
- AF3 max_ranking_score and avg_ranking_score measure predicted-structure
  confidence, not the probability that a biological stoichiometry is correct.
- Compare both maximum and average scores. A high maximum with a much lower
```

```
  average may indicate that confidence varies among the generated models.
- Consider num_models when comparing maximum or average scores; scores based
  on different numbers of predictions are not necessarily directly comparable.
- Missing AF3 scores mean unavailable structural evidence, NOT a score of zero
  or negative biological evidence, including when a complex is too large to model.

{{ASSEMBLY_SIZE_INTERPRETATION_RULES}}
```

**AlphaFold3 rules for homomeric targets.**

```
ALPHAFOLD3 EVIDENCE RULES:
- Only max_ranking_score is supplied as AF3 evidence for this target.
- This is predicted-structure confidence, not the probability that a biological
  stoichiometry is correct. Weigh it with the supplied Stage-1 support and
  biological evidence; it does not automatically determine the final ranking.
- Missing AF3 scores mean unavailable structural evidence, NOT a score of zero
  or negative biological evidence, including when a complex is too large to model.
- Do not infer unprovided AF3 statistics, structural observations, or consistency
  across predictions from the maximum score alone.

{{ASSEMBLY_SIZE_INTERPRETATION_RULES}}
```

**Assembly-size rules appended to either AlphaFold3 block.**

```
EXPLICIT ASSEMBLY-SIZE INTERPRETATION RULES:
- Compare AF3 scores cautiously when candidates differ in total residue count,
  number of chains, or copy numbers (for example, A1B1 versus A2B2). A larger
  assembly introduces additional relative chain arrangements and potential
  interfaces, which can reduce global confidence even when the larger
  stoichiometry is biologically correct.
- A higher AF3 ranking score for a smaller complex is NOT, by itself, evidence
  that the smaller stoichiometry is biologically correct. Do not automatically
  prefer the smaller candidate merely because its AF3 score is higher.
- Do not automatically prefer a larger candidate merely because it has more
  copies, greater symmetry, or more apparently occupied binding sites.
- Look for target-specific biological evidence supporting or contradicting the
  additional subunit copies and interactions required by a larger candidate.
  If only global AF3 scores are supplied, do NOT invent interface-level scores,
  contacts, clashes, or structural observations that have not been provided.
- Do NOT apply any fixed correction, bonus, penalty, or numerical normalization
  of AF3 scores by sequence length, total residues, number of chains, or copies.
  Instead, assess size-related uncertainty when interpreting the raw scores.
- Neither the highest AF3 score nor the highest Stage-1 support must win.
  If the structural scores and biological evidence disagree, briefly explain
  the tradeoff and any uncertainty behind the final ordering.
```

**Evidence rules for the no-AlphaFold3 control.**

These replace the AlphaFold3 rules when that control is run.

```
EVIDENCE RULES:
- Rank using the target description, sequences, and Stage-1 support information.
- Stage-1 rankings and support values are fallible and are not probabilities.
- Do not automatically favor the highest Stage-1 support or simplest stoichiometry.
- Do not assume knowledge of AlphaFold3 results or other unseen evidence.
```

**Stage-2 user message.**

```
Target information (do not infer a benchmark ID):
```

```
Description: {{DESCRIPTION_OR_NOT_AVAILABLE}}
Organism/source: {{ORGANISM_OR_NOT_AVAILABLE}}
Subunit sequences (each letter identifies a distinct protein):
{{SUBUNIT_ID}} ({{LENGTH}} residues): {{AMINO_ACID_SEQUENCE}}
{{REPEAT_FOR_EACH_UNIQUE_SUBUNIT}}

{{OPTIONAL_STAGE_1_RESEARCH_TEMPLATE_EVIDENCE_BLOCK}}

Stage-1 candidate pool (ONLY these candidates are allowed):
- {{STOICHIOMETRY_1}}: Stage-1 rank={{RANK_1}}; Stage-1 support={{SUPPORT_1}}{{
    OPTIONAL_SUPPORT_EVIDENCE_1}}{{CONDITION_SPECIFIC_AF3_FIELDS_1}}
{{REPEAT_FOR_EACH_SHORTLISTED_CANDIDATE_IN_STAGE_1_ORDER}}

Use precisely this PLAIN-TEXT output format (not JSON):
Candidates:
1. {{STOICHIOMETRY_1}}
2. {{STOICHIOMETRY_2}}
{{CONTINUE_THROUGH_ALL_SUPPLIED_CANDIDATES}}
Final Answer: {{STOICHIOMETRY_1}}
Support Evidence: A concise explanation of the most important evidence and uncertainty.
The example order above is only the ORIGINAL Stage-1 order; independently decide how to re-
    rank it.
Return every input candidate exactly once; list the ranking and Final Answer FIRST.
Do not generate another numbered list in Support Evidence; prose there is free-form.
Your ranking is valid even if the explanation is short; prioritize completing the ranking.
```
